# Resident microbial communities inhibit growth and antibiotic resistance evolution of *Escherichia coli* in human gut microbiome samples

**DOI:** 10.1101/741439

**Authors:** Michael Baumgartner, Florian Bayer, Katia R. Pfrunder-Cardozo, Angus Buckling, Alex R. Hall

## Abstract

Countering the rise of antibiotic resistant pathogens requires improved understanding of how resistance emerges and spreads in individual species, which are often embedded in complex microbial communities such as the human gut microbiome. Interactions with other microorganisms in such communities might suppress growth and resistance evolution of individual species (e.g. via resource competition), but could also potentially accelerate resistance evolution via horizontal transfer of resistance genes. It remains unclear how these different effects balance out, partly because it is difficult to observe them directly. Here, we used a gut microcosm approach to quantify the effect of three human gut microbiome communities on growth and resistance evolution of a focal strain of *Escherichia coli*. We found the resident microbial communities not only suppressed growth and colonization by focal *E. coli*, they also prevented it from evolving antibiotic resistance upon exposure to a beta-lactam antibiotic. With samples from all three human donors, our focal *E. coli* strain only evolved antibiotic resistance in the absence of the resident microbial community, even though we found resistance genes, including a highly effective resistance plasmid, in resident microbial communities. We identified physical constraints on plasmid transfer that can explain why our focal strain failed to acquire some of these beneficial resistance genes, and we found some chromosomal resistance mutations were only beneficial in the absence of the resident microbiota. This suggests, depending on *in situ* gene transfer dynamics, interactions with resident microbiota can inhibit antibiotic resistance evolution of individual species.

## Introduction

The over- and inappropriate use of antibiotics has promoted the evolution of resistance in pathogens, resulting in a crisis for human health care [1]. To combat this problem, it is important to understand the underlying mechanisms of how resistance is acquired by bacteria and spreads within bacterial populations and communities [2, 3]. A large body of research has used direct observations of resistance evolution in simplified laboratory conditions to understand how antibiotics drive the spread of resistance [4, 5]. A key limitation of this approach is that it excludes interactions with other microorganisms, which we can expect to be important for bacteria evolving in natural or clinical settings because they spend most of their time in dense and diverse microbial communities. Interactions in species-rich microbial communities might negatively affect growth of individual species via, for example, competition for resources or niche space [6]. This may in turn inhibit antibiotic resistance evolution of individual species, because reduced population growth should reduce the supply of new genetic variation. On the other hand, interspecific interactions also potentially have positive effects on growth and evolution of individual species via, for example, exchange of genetic material [7], cross-feeding or public goods sharing [9–11]. Community-level interactions can also alter the strength of selection for resistant variants in the population [12, 13]. In support of a key role for interspecific interactions in resistance evolution, observations of bacteria isolated from natural and clinical settings indicate genes involved in antibiotic resistance evolution are often horizontally transferable [14–17]. Despite this, direct observations of how these different types of effects balance out are lacking. Consequently, it remains unclear how interactions with species-rich microbial communities affect growth and antibiotic resistance evolution of individual species or strains of bacteria.

The impact of interactions with other microorganisms for antibiotic resistance evolution is likely to be particularly important in the human gastrointestinal tract. This is one of the most densely inhabited environments in the world, colonized by a rich diversity of bacteria, viruses and eukarya, which are embedded in a network of biotic interactions [18, 19]. Interactions among microorganisms in the gut microbiome (which we take here to mean the resident microorganisms, their genes and the local abiotic environment, following Marchesi & Ravel [20] and Foster et al. [19]) play an important role for human health [21]. For example, the microbiome minimizes potential niche space for invading species, making it harder for them to establish in the community, thereby contributing to colonization resistance against pathogens [22, 23]. This suggests competitive interactions with other microorganisms are common, which we would expect to inhibit population growth of individual taxa and in turn constrain their ability to evolve antibiotic resistance. On the other hand, some interactions in the gut may be mutualistic [10] or modify the effects of antibiotics on individual species [24], potentially resulting in a net positive effect on growth. Moreover, recent metagenomic studies [25–27] showed the gut microbiome harbours a variety of mobile genetic elements, often carrying resistance and virulence genes, that are shared by community members. Consistent with this, horizontal transfer of resistance genes within individual hosts is central to resistance evolution in several key pathogens found in the gastrointestinal tract [28–31]. This suggests interactions with other microorganisms in the gut microbiome can also promote growth and resistance of individual taxa. We aimed to quantify the net effect of interactions with species-rich communities of other microorganisms, in particular those found in the human gastrointestinal tract, for growth and resistance evolution of a given strain that newly arrives in the community.

We approached this question using a human gut microcosm system consisting of anaerobic fermentors filled with human faecal slurry, including the resident microbial community and the beta-lactam antibiotic ampicillin, to which bacteria can evolve resistance by chromosomal mutations [32] or horizontal acquisition of beta-lactamase genes [33]. We used ampicillin because beta-lactam antibiotics are very widely used in human healthcare [34], resistance is a major problem [35], and key mechanisms by which bacteria evolve resistance to ampicillin overlap with resistance mechanisms against other antibiotics [36]. Because the microbiota in faecal samples reflects the diversity of the distant human gastrointestinal tract [37], this approach allowed us to produce microcosms containing species-rich communities sampled from human gut microbiomes. We aimed to determine how interactions with this resident microbial community affected growth and resistance evolution of *Escherichia coli*. We focused on *E. coli* because it is a ubiquitous gut commensal [38] and key opportunistic pathogen [39] for which antibiotic resistance is an increasing problem [40]. We inoculated each microcosm with a tagged, focal *E. coli* strain, before tracking its growth and resistance evolution in the presence and absence of ampicillin. By also including microcosms containing sterilized versions of the same faecal slurry (where the resident microbial community had been deactivated), we quantified the net effect of interactions with the resident microbial community. This approach allowed us to (i) track growth and resistance evolution of the focal strain in the presence and absence of resident microbial communities sampled from several human donors; (ii) isolate plasmid-carrying *E. coli* strains from the resident microbial community and identify constraints on horizontal transfer of resistance genes; (iii) characterize the resident microbial communities and how they changed over time. Our results show the resident microbial community inhibits both growth and resistance evolution of *E. coli*, despite the presence of resistance plasmids that can be conjugatively transferred to our focal strain in certain physical conditions.

## Results

### Resident microbial communities suppressed growth of a focal *E. coli* strain

We cultivated our focal *E. coli* strain in anaerobic microcosms in the presence and absence of an antibiotic and three different samples of gastrointestinal microbiomes, each from a different human donor, for seven days (S1 Fig). On average, antibiotic treatment (ampicillin) decreased focal strain abundance (effect of antibiotic in a generalized linear mixed model with zero inflation, glmmadmb, *χ*^2^ = 14.14, d.f. = 1, *P* < 0.001; Fig 1). For example, after 24h focal strain abundance was reduced compared to ampicillin-free treatments by 69% (s.d.=6.02) in the basal medium treatment, 78-90% (depending on human donor) in the sterilized slurry treatments, and 84-99.9% (depending on human donor) in the “live” slurry treatments (Fig 1; S1 Table). Inclusion of the resident microbial community from human faecal samples also reduced focal strain abundance on average, which we inferred by comparing the community treatments (“live” faecal slurries, including the resident microbial community) to the community-free treatments (sterilized versions of the same faecal slurries; effect of community in glmmadmb, *χ*^2^ = 6.65, d.f. = 1, *P* = 0.01; Fig 1).

**Fig 1.**
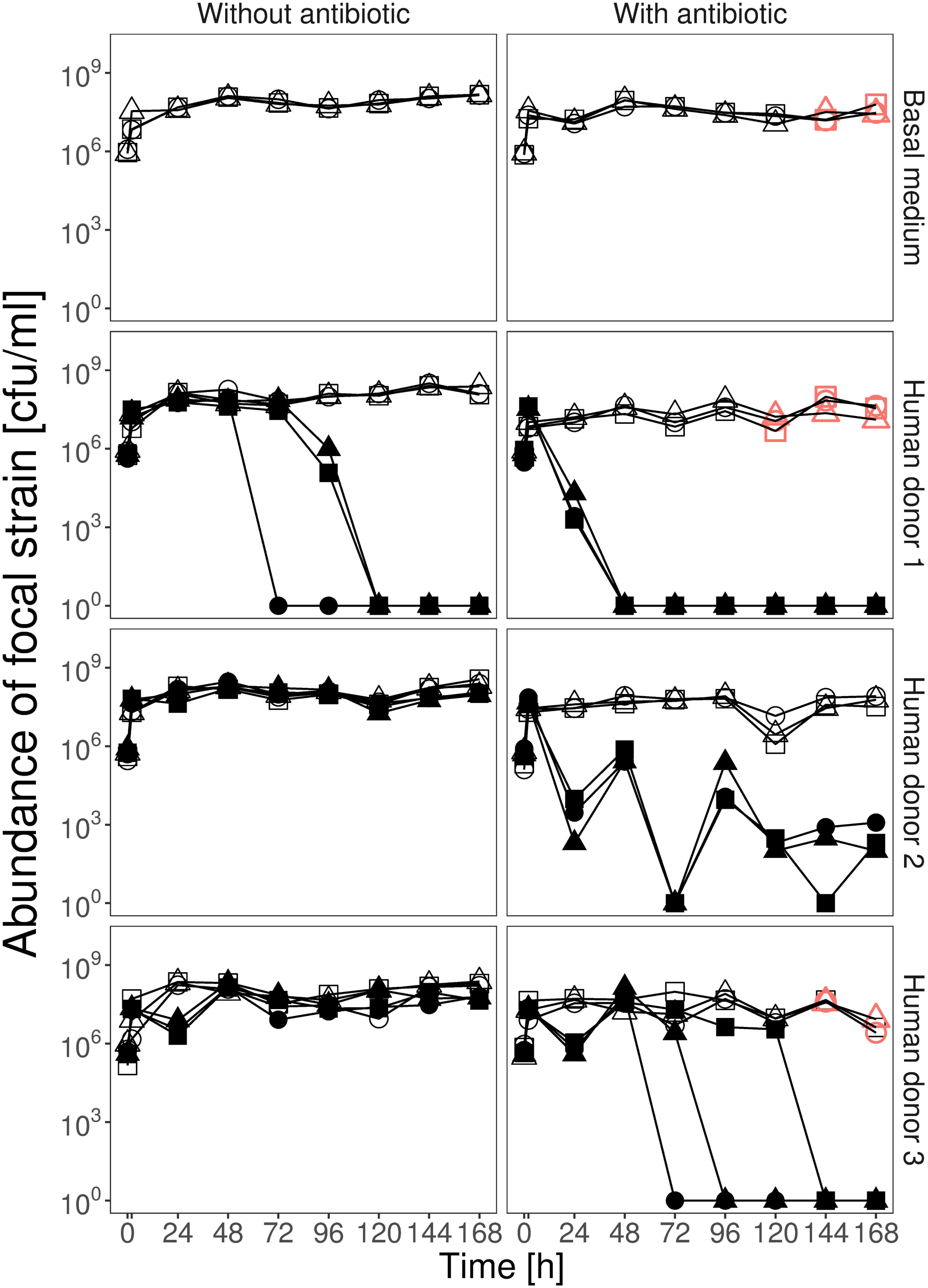
Resident microbial communities suppressed growth and resistance evolution of focal *E. coli*. Each panel shows abundance of the focal *E. coli* strain (in colony-forming units per ml) over seven days for either basal medium (top row) or with faecal slurry from one of three human donors, in the absence (left panels) or presence (right panels) of ampicillin, which was applied at the sampling time points after 2h and thereafter at each daily transfer. Empty symbols show community-free treatments; filled symbols show treatments with the resident microbial community; red symbols show microcosms where we detected ampicillin-resistant colony isolates of the focal strain. The three lines, each with different symbols, in each treatment show three replicate microcosms. Microcosms where we detected no focal strain colonies are shown at 10^0^.

The suppressive effect of resident microbial communities depended on both which human donor sample was used to prepare the microcosms (donor×community interaction in glmmadmb, *χ*^2^ = 10.23, d.f. = 2, *P* = 0.006) and the presence of ampicillin (antibiotic×community interaction in glmmadmb, *χ*^2^ = 5.2, d.f. = 1, *P* = 0.02), being strongest for populations exposed to both resident microbiota and the antibiotic (Fig 1). This resulted in extinction of the focal strain (below our detection limit) in populations exposed to ampicillin and the resident microbial communities from human donors 1 and 3. That is, in these treatments the focal strain failed to colonize the community. For the resident community from human donor 2, the focal strain was driven to very low abundance in the presence of the community and ampicillin together, but did not disappear completely (Fig 1). In the absence of ampicillin, resident communities still suppressed the focal strain on average (effect of community in glmmadmb for ampicillin-free treatments only, *χ*^2^ = 10.04, d.f. = 1, *P* = 0.002). As in the presence of ampicillin, the strength of this effect varied depending on human donor, being strongest for the resident microbial community from human donor 1 (effect of community for this donor in the absence of ampicillin: glmmadmb, *χ*^2^ = 11.28, d.f. = 1, *P* = 0.0002; Fig 1; S1 Table), resulting in exclusion of the focal strain. The resident community from human donor 3 suppressed average growth of the focal strain by ∼54% across the entire experiment (effect of community for human donor 3 in the absence of ampicillin: glmmadmb, *χ*^2^ = 4.77, d.f. = 1, *P* = 0.03; Fig 1; S1 Table). For the resident community from human donor 2, average focal strain abundance was lower in the presence of the community (mean reduction of 24% compared to community-free microcosms; Fig 1; S1 Table), although this was not statistically significant (effect of community for human donor 2 in the absence of ampicillin: glmmadmb, *χ*^2^ = 0.66, d.f. = 1, *P* = 0.41). We found no evidence that abiotic factors in the sterile faecal slurry were suppressive for the focal strain: there was no significant variation in average focal strain abundance among the community-free treatments and the control treatment containing only the basal growth medium that was used to prepare faecal slurries (linear mixed effects model, glmer: *χ*^2^ = 0.41, d.f. = 3, *P* = 0.94). In summary, the resident microbial communities sampled from three human donors each suppressed growth of a focal *E. coli* strain in anaerobic fermentors filled with faecal slurry, but to varying degrees, and this effect was amplified by adding ampicillin.

### Stable total bacterial abundance but variable community composition over time

We used flow cytometry to estimate total bacterial abundance in microcosms containing resident microbial communities. In antibiotic-free microcosms, total bacterial abundance was approximately stable over time (>10^9^ cells/ml; Fig 2), and was higher on average than in microcosms exposed to ampicillin (effect of antibiotic in a linear mixed effects model, lmm: *χ*^2^ = 10.37, d.f. = 1, *P* = 0.001). However, the suppressive effect of the antibiotic varied over time (antibiotic×time interaction in lmm: *χ*^2^ = 101.81, d.f. = 7, *P* < 0.001), being strongest at the beginning of the experiment. The effect of the antibiotic also varied across communities from different human donors (antibiotic×donor interaction in lmm: *χ*^2^ = 79.30, d.f. = 2, *P* < 0.001), with those from human donors 1 and 2 showing a stronger recovery after the first application of ampicillin (which resulted in a drop in abundance after 24h) than the community from human donor 3. These results show our experimental set-up sustained high numbers of microorganisms in the community treatments over time in both the presence and absence of ampicillin.

**Fig 2.**
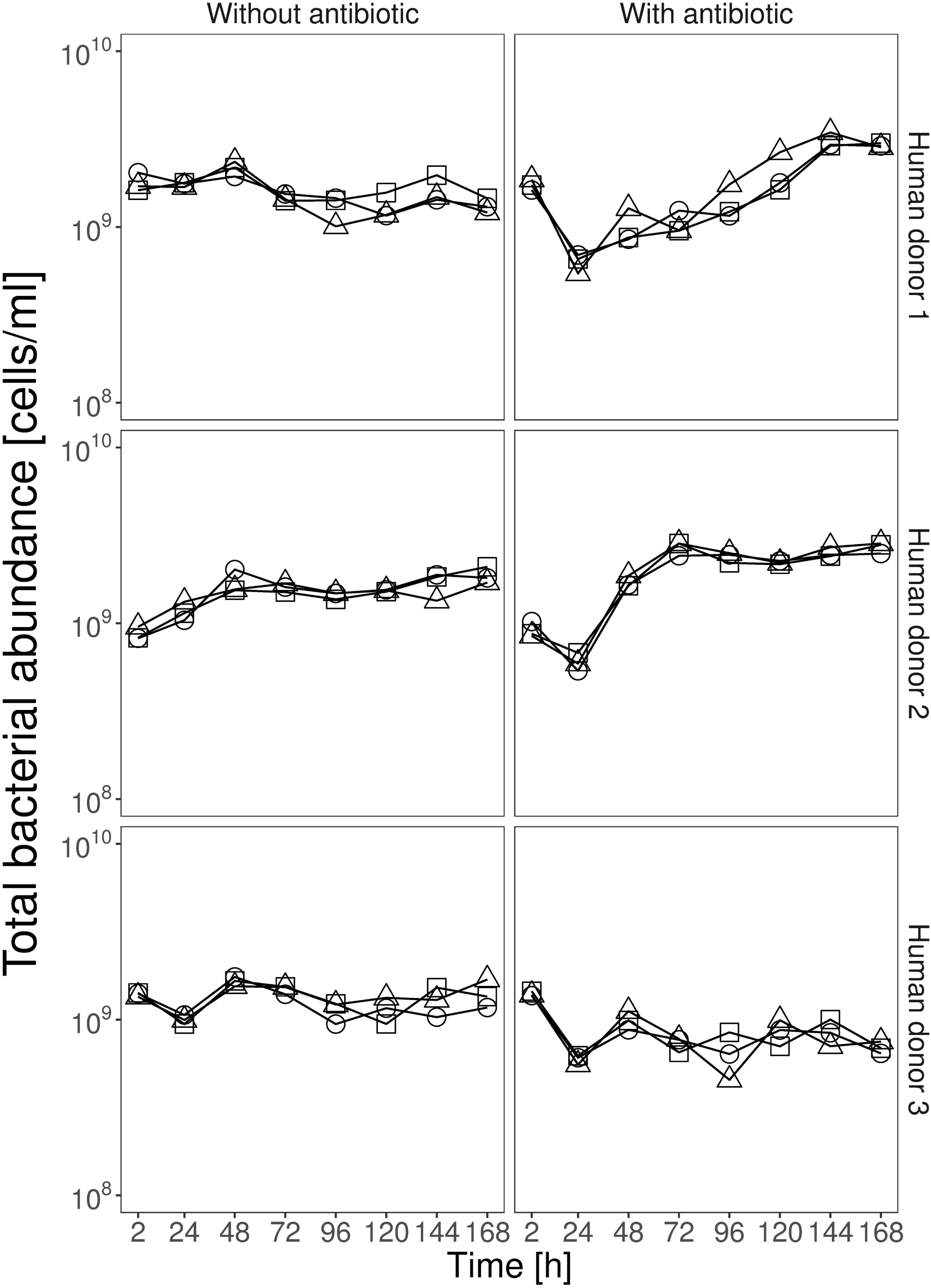
Total bacterial abundance over time in community treatments with and without antibiotics. Each row of panels shows data from one of the three human donors, and the right/left panels show treatments with/without ampicillin. The three replicate communities in each panel are shown by three different lines, each with a different symbol. Total bacterial abundance was measured by flow cytometry (see Methods).

To investigate the community composition in these microcosms, we used amplicon sequencing of the variable regions 3 and 4 of the 16S rRNA gene. This revealed similar levels of within-sample diversity (Shannon’s alpha diversity; Fig 3A) in microbiome samples from the three human donors at the start of the experiment. Within-sample diversity then decreased slightly over the first 24h of the experiment, and significantly between 24h and 168h (effect of time in a linear mixed effects model including data from 24h and 168h, lme: *F* _1, 14_ = 481.43, *P* < 0.001). This applied across the three human donors (effect of human donor, lme: *F* _2, 15_ = 2.07, *P* = 0.16), which also showed similar shifts in taxonomic composition over time (Fig 3B). Communities at 0h were dominated by the families *Lachnospiraceae* and *Ruminococcaceae*, plus *Prevotellaceae* for human donor 3. Over time these groups became less abundant relative to *Enterobacteriaceae* and *Bacteroidaceae*. Analysis of the reads assigned to *Enterobacteriaceae* indicated *E. coli* was always the most abundant member of this group (see Methods), accounting for ∼99% of the 16S rRNA amplicon reads assigned to *Enterobacteriaceae* in all samples (S2 Table), with the remaining ∼1% comprising other species (e.g. *Enterobacter sp., Citrobacter sp.* and *Klebsiella sp.*). Compared to changes over time, ampicillin had a weak effect on within-sample diversity (effect of antibiotic: *F* _1, 14_ = 11.37, *P* = 0.006). This interpretation was supported by an alternative analysis (Principal Coordinate Analysis; S2 Fig) based on Bray-Curtis dissimilarities. Thus, despite approximately stable total abundance, we saw changes in community composition over time that were more pronounced than differences among communities from different human donors or antibiotic treatments. Despite these changes in relative abundance of different taxa, the identities of the top 5-6 families were stable over time and across human donors (Fig 3B).

**Fig 3.**
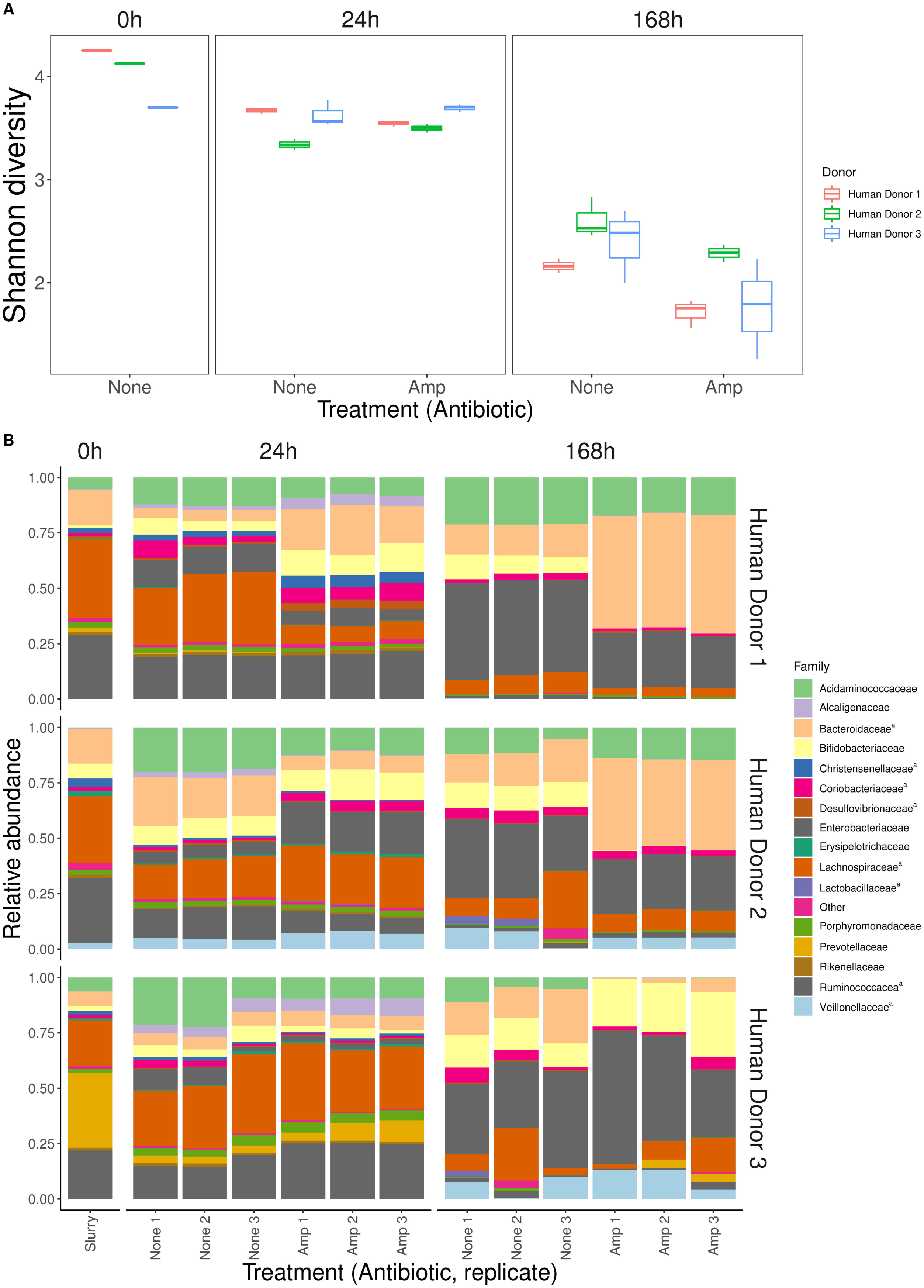
Within-sample diversity and changes in community composition over time. **(A)** Diversity is estimated here using Shannon’s diversity index for samples from three timepoints (0h, 24h, 168h, shown at top) and three human donors (see legend), in the presence and absence of ampicillin (*x*-axis). Each box shows data from three replicate microcosms per treatment group and time point (except for at 0h, which shows the single initial sample from each human donor). **(B)** Relative abundance of the 15 most prevalent bacterial families in each microcosm at three timepoints (0h, 24h and 168h, shown at top), for each human donor (rows of panels, labelled at right) in the presence and absence of ampicillin (three replicates each treatment; *x*-axis). Superscript “^a^” in the legend indicates obligately anaerobic families.

We used quantitative PCR to better understand the contribution of resident *E. coli* and the focal strain to the total abundance of *E. coli* (see S1 Methods). Consistent with the amplicon data above, this revealed increasing total abundance of *E. coli* sequences over time in both the presence and absence of ampicillin (S3 Fig). The copy number of focal strain sequences relative to total *E. coli* indicated the focal strain was rare relative to other *E. coli* after 24h (S3 Fig; S2 Table). At the end of the experiment, consistent with our estimates above from selective plating and colony PCR, the focal strain was below the detection limit in treatments containing the community from human donor 1, both with and without ampicillin, and the human donor 3 community with ampicillin; in the other treatments focal strain sequences were rare compared to total *E. coli* (S2 Table).

### Antibiotic resistance evolved only in the absence of resident microbial communities

We screened for the emergence of antibiotic-resistant variants of the focal strain (that had acquired resistance to ampicillin) after every growth cycle by plating each population onto antibiotic-selective plates (8 μg/ml ampicillin; approximating the minimal inhibitory concentration (MIC) of the focal strain). We never observed resistant variants of the focal strain in any of the community treatments (populations exposed to the resident microbial communities from human microbiome samples). By contrast, in community-free treatments (basal growth medium and sterilized human faecal slurries) resistant variants appeared toward the end of the experiment at 120h (slurry from human donor 1) and 144h (basal growth medium and slurry from human donor 3), although not in sterilized samples from human donor 2 (Fig 1). Thus, the resident microbial community from human microbiome samples suppressed antibiotic resistance evolution in our focal strain.

To investigate genetic mechanisms associated with resistance evolution and general adaptation to our experimental conditions, we performed whole-genome sequencing for two sets of focal strain isolates from the final time point: 8 ampicillin-resistant colony isolates from ampicillin plates (one from each of the eight populations where we observed the emergence of antibiotic resistance during the experiment) and 33 randomly selected colony isolates from ampicillin-free plates (each from a different population and across all treatments). In the antibiotic-resistant isolates, all SNPs and the deletion we found (Table 1) were in genes related to membranes (*ompR, ftsI, opgB*), general stress response (*rpoC, relA*) or transcription regulation (*rpoD*). Two genes were mutated independently in multiple colony isolates: *rpoC* and *rpoD*. We also detected an insertion sequence (IS) movement between *perR* and *insN* in two colony isolates. Of genes where we detected mutations in a single colony isolate, *ftsI* [41], *relA* [42] and *ompR* [43] have each been previously annotated as being involved in resistance to beta-lactam antibiotics. *ompR* was also mutated in three randomly selected colony isolates, all from populations that had been exposed to ampicillin during the experiment (S3 Table). We found six other genes mutated in parallel in between 2 and 5 randomly selected colony isolates (S3 Table). Five of these were mutated in isolates from both antibiotic and antibiotic-free treatments. This included *insN* (involved in transposition) and *gtrS* (involved in aerobic respiration), both mutated in five isolates. Across the two sets of colony isolates (randomly selected and antibiotic-resistant), three other loci were mutated in both sets. These were *rpoD* (only in isolates that had been exposed to antibiotics), *opgB* and *yaiO* (both in isolates from antibiotic and antibiotic-free treatments). In summary, we found some parallel genetic changes specific to antibiotic treatments and consistent with known resistance mechanisms, plus other genetic changes that occurred across antibiotic and antibiotic-free treatments and are therefore more likely involved in general adaptation.

**Table 1:** Genes mutated in ampicillin-resistant colony isolates of the focal strain from the end of the experiment. Asterisks indicate genes also mutated in randomly selected clones from ampicillin-free plates (see S3 Table).

| Treatment group | Human donor | Replicate | *afuB* | *afuC* | *cspB* ⇄ *cspF* | *cyoA* | *fecI* | *ftsI* | *insA* ⇄ *uspC* | *insB1* | *insD1* | *ompR* | *opgB* * | *perR* | *perR* ⇄ *insN* | *relA* | *rpoC* | *rpoD* | *yaiO* * | *yehE* | *yjhD* ⇄ *insO* | *yjhD* ⇄ *yjhE* |
| --- | --- | --- | --- | --- | --- | --- | --- | --- | --- | --- | --- | --- | --- | --- | --- | --- | --- | --- | --- | --- | --- | --- |
| Basal medium with antibiotic | None | 1 |  |  | x |  |  | ● |  |  |  |  |  |  | x |  |  |  |  |  |  | x |
|  |  | 2 |  | x |  |  |  |  |  |  |  |  |  |  |  |  |  |  |  | x |  |  |
|  |  | 3 |  |  |  |  |  |  |  |  |  |  |  |  | x | ● | ● |  |  |  |  |  |
| Community-free with antibiotic | 1 | 1 |  |  |  |  |  |  | x |  | x |  |  | x |  |  |  | ● |  |  |  |  |
|  |  | 2 |  |  |  |  |  |  |  |  |  | ● |  |  |  |  |  |  |  |  |  |  |
|  |  | 3 | x |  |  | x |  |  |  |  |  |  | ● |  |  |  |  |  | x |  |  |  |
| Community-free with antibiotic | 3 | 1 |  |  |  |  |  |  |  | x |  |  |  |  |  |  |  | ● |  |  |  |  |
|  |  | 2 |  |  |  |  | x |  |  |  |  |  |  |  |  |  | ▲ |  |  |  | x |  |

▲ Deletion x Insertion ● SNP

### Plasmid acquisition was constrained by lack of transfer, not lack of fitness benefits

We next sought to explain why we never observed antibiotic resistance evolution of the focal strain in the presence of resident microbial communities, which we had expected to harbour beneficial resistance genes [26,27,44–46]. We hypothesized this could have been due to a lack of horizontally transferable resistance genes in the resident microbial communities. However, we detected ampicillin resistant *E. coli* in the resident microbial communities from human donors 1 and 3 (by selective plating), and after sequencing their genomes we identified several antibiotic resistance genes that were associated with plasmid genes (Fig 4; S4 Table). In the hybrid assembly (using MinION and Illumina reads) of a representative isolate from human donor 1, we identified two plasmids. The larger plasmid had four known resistance genes (Fig 4A), including one conferring resistance to beta-lactam antibiotics. We also identified three IncF replicons on this plasmid and a complete set of *tra* genes, which are involved in conjugative plasmid transfer. The second plasmid carried a known replicon (ColRNAI), and mobilization genes (*mbeA* and *mbeC*), plus a complete colicin E1 operon (*cnl, imm, cea*). For a representative isolate from human donor 3, we found the plasmid replicon and resistance genes integrated on the chromosome (Fig 4B). This putative integrated plasmid from human donor 3 also carried multiple resistance genes, including a beta-lactamase and an IncQ replicon, which is a part of the *repA* gene [47], but we detected no *tra* genes. The other five genome assemblies (Illumina reads for other isolates from the same human donor) contained the same resistance genes and replicons across multiple smaller contigs (S4 Table). Mapping the corresponding sequencing reads of all Illumina-sequenced isolates to the long-read data single contig found in the isolates sequenced on the MinION platform revealed identical mapping in all ten cases, consistent with these genomes having the same structure across each of the isolates (S4 Table).

**Fig 4.**
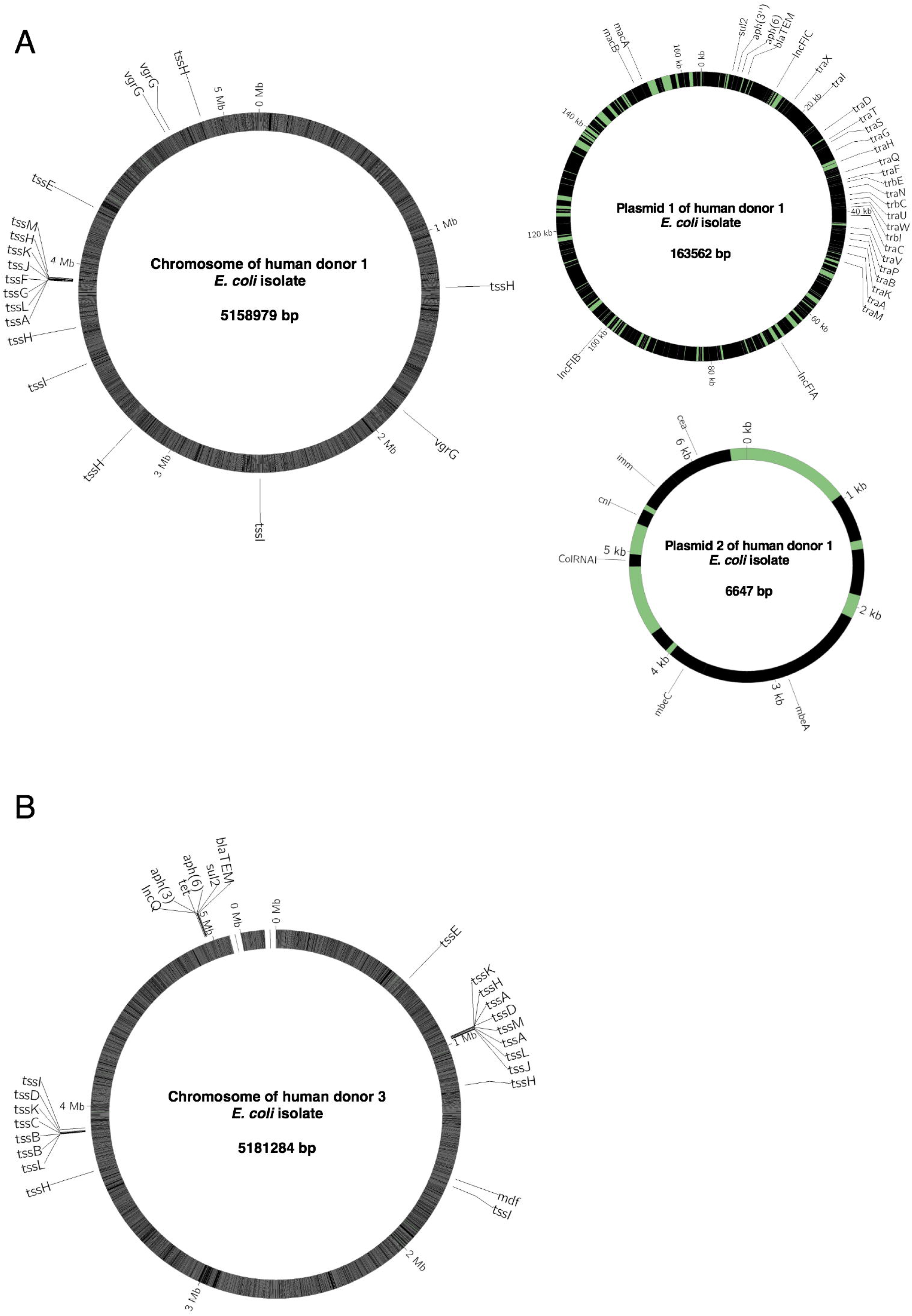
Schematic maps of plasmids and chromosomes for representative resident *E. coli* isolates from (A) human donor 1 and (B) human donor 3. Sequences are annotated with known plasmid replicon sequences (IncFIA, IncFIB, IncFIC on plasmid 1 and ColRNAI on plasmid 2 in (A); IncQ on the chromosome in (B)), genes involved in horizontal transfer (*tra* and *trb*), known resistance genes (*bla_TEM_*, *sul2*, *aph(3), aph(6), mdf*), genes involved in Type 6 secretion systems (*tss, vgr*), genes involved in colicin production and immunity (*cea, cnl, imm*) and mobilization (*mbeA* and *mbeC*). The genome of the isolate from human donor 3 (B) is not closed, as indicated with a gap. Colours indicate coding (black) and non-coding regions (green); note the scale varies among chromosomes and plasmids.

We hypothesized that the lack of plasmid-driven resistance evolution in our focal strain might have been caused by constraints on conjugative transfer that made these plasmids inaccessible. Using a conjugative mating assay on agar, we never found transconjugants of our focal strain when it was mixed with an isolate from human donor 3 (identified above as carrying a putative integrated plasmid). This is consistent with the lack of *tra* genes on this plasmid and suggests it could not be transferred into our focal strain by conjugation in the absence of other drivers of horizontal gene transfer (e.g. phages or other plasmids). This is also consistent with past work suggesting IncQ plasmids are mobilizable rather than conjugative [48, 49], and that we did not detect any other plasmid replicons in the same isolates. However, for the plasmid from human donor 1 we found transconjugants of our focal strain at the end of the mating assay, which we confirmed by colony PCR (S4 Fig). This suggests this plasmid was conjugative and could be transferred to our focal strain, consistent with the presence of *tra* genes on this plasmid (Fig 4A).

Given that the resistance plasmid from human donor 1 was transferable into our focal strain, why did it not spread in the main experiment above? We hypothesized this could result from plasmid-borne resistance being less beneficial than resistance acquired by chromosomal mutation (as we observed in community-free treatments in the main experiment). We would expect a net benefit of resistance to result in increased population growth at the antibiotic concentration applied during the experiment (7.2 μg/ml). We found acquisition of the plasmid conferred a much larger increase in population growth across all non-zero antibiotic concentrations than that observed for evolved colony isolates that had acquired resistance via chromosomal mutation during the main experiment (Fig 5A; S5 Table). Furthermore, in pairwise competition experiments, the transconjugant carrying this plasmid had a strong competitive advantage relative to the wild type in the presence of the resident microbial community from human donor 1 (S5 Fig A). This fitness advantage was increased by adding ampicillin at the concentration we used in the main experiment, and even further by adding ampicillin at three times the IC90 of the ancestral focal strain (community × ampicillin interaction by permutation test; *P* = 0.029; S5 Fig B). This shows it would have been highly beneficial for the focal strain to acquire the plasmid in our experiment, particularly in the presence of ampicillin.

**Fig 5.**
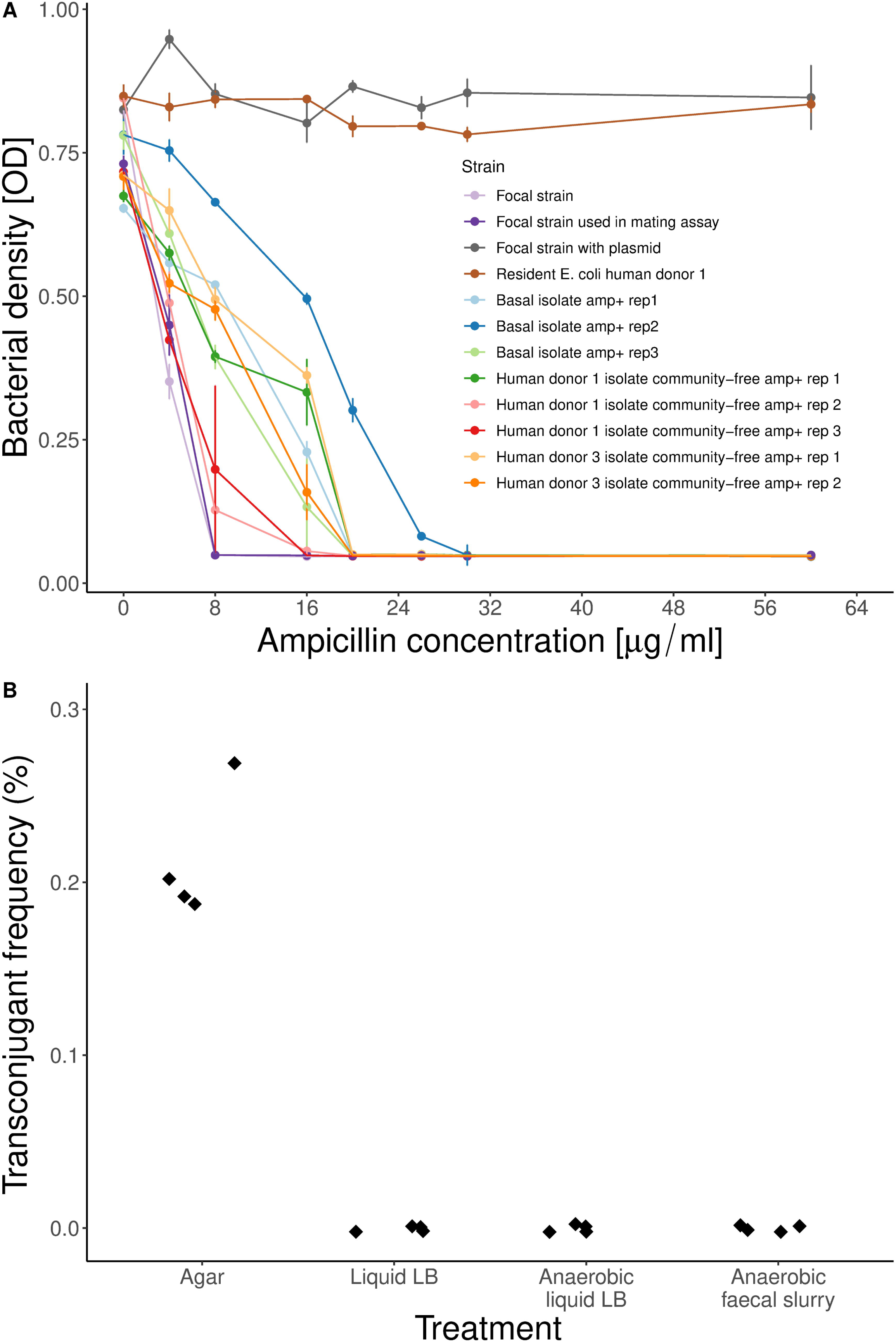
Transfer of a resistance plasmid from the resident microbial community is sensitive to abiotic conditions, and it confers a large increase in resistance. **(A)** Antibiotic susceptibility of the ancestral focal strain, the version of the focal strain used to isolate transconjugants (see Methods), the focal strain with the plasmid (transconjugant), the resident *E. coli* isolate used as the plasmid donor, and eight evolved focal strain colony isolates that we isolated on ampicillin plates and that had chromosomal mutations (S2 Table). Average optical density (OD) values ±s.e. are shown after 24h growth at each ampicillin concentration. **(B)** Transconjugant frequency (as a percentage of the total recipient population) after mating experiments in various conditions. The recipient strain was a tagged version of the ancestral focal strain and the plasmid donor was a resident *E. coli* isolate from human donor 1. For the faecal slurry treatment we used sterilized faecal slurry from human donor 1.

Unlike transconjugants carrying the plasmid from resident *E. coli*, two evolved colony isolates of the focal strain carrying chromosomal resistance mutations had a fitness advantage relative to the wild type only in the absence of the community; in the presence of the resident microbial community they had a fitness disadvantage (effect of community by permutation test: *P <* 0.001 for isolates from human donors 1 and 3; S5 Fig B). This suggests the genetic changes associated with increased resistance in these isolates in the absence of resident microbiota would not have been beneficial in the Community+Ampicillin treatments of the main experiment, unlike the plasmid we isolated from resident *E. coli*. This conclusion was supported by direct competitions between transconjugants carrying the resistance plasmid and evolved colony isolates from human donors 1 and 3 (S6 Fig). Further competition experiments showed the competitive advantage of resident *E. coli* from human donors 1 and 3 carrying resistance genes was also present in the absence of other resident microbiota (S7 Fig).

Another possible explanation for the lack of transfer of the resistance plasmid from human donor 1 in the main experiment is that conjugative transfer might be specific to particular environmental conditions. This has been observed for other plasmids across various experimental conditions [50, 51]. We tested this by mating assays as above, but in a range of different experimental conditions. We detected transconjugants that had acquired the plasmid at a final frequency of ∼0.2% (as a fraction of the total recipient population) after mixing the plasmid-carrying isolate from human donor 1 and the focal strain on an agar surface, but we found no transconjugants after doing the same experiment in three different types of liquid growth medium (lysogeny broth (LB), anaerobic LB, and anaerobic community-free faecal slurry; Fig 5B). This shows transfer of the conjugative plasmid we isolated from human donor 1 requires particular abiotic conditions, which may explain why our focal strain failed to evolve resistance via horizontal gene transfer in the presence of resident microbial communities. This was supported by simulations of a hypothetical plasmid with similar properties but that is transferable in our gut microcosm system (S1 Model). This indicated that, if the plasmid from human donor 1 had been conjugatively transferable in our gut microcosm system, we would have detected transconjugants in our main experiment (although only with relatively high transfer rates; S1 Model). The same model also showed growth suppression of invading lineages by resident microbiota can reduce transconjugant abundance, suggesting even when horizontal acquisition of beneficial resistance genes is common, interactions with resident microbiota can impede their spread.

## Discussion

We found resistance via chromosomal mutation to an important class of antibiotics (beta-lactams) evolved in a focal *E. coli* strain in our experiment only in the absence of resident microbial communities sampled from healthy human volunteers. The suppressive effect of these resident microbial communities was strong enough that the focal *E. coli* strain was driven towards extinction (below our detection limit) when it was exposed to both ampicillin and the community simultaneously (with communities from two of the three human donors we tested). Consequently, the net effect of the resident microbial communities here was to confer a form of colonization resistance against a non-resident strain, and to prevent that strain from evolving antibiotic resistance. Our analysis of resident *E. coli* isolates (not the focal strain) from the microbial communities showed this occurred despite the presence of beneficial, potentially horizontally transferable resistance genes. Genomic analyses and conjugation experiments with these resident *E. coli* isolates showed the *in situ* transfer dynamics depend critically on genetic (the presence of genes encoding the machinery for conjugative transfer) and abiotic (physical structure of the environment) factors. This is important because ultimately it is the *in situ* transfer dynamics that will determine whether or not horizontal transfer of beneficial resistance genes is sufficient to counteract the growth-suppressive effects of interactions with the community and confer a net benefit to invading lineages. Community-level interactions also modified selection for resistance, amplifying growth inhibition by ampicillin and altering the relative advantages/disadvantages of individual resistant genotypes (evolved isolates with chromosomal mutations conferring relatively weak increases in ampicillin resistance had a reduced advantage, but plasmids from resident *E. coli* that conferred relatively large increases in ampicillin resistance were more beneficial in the presence of resident microbiota). Overall, this indicates resident microbiota influence resistance evolution of invading strains via effects on both population dynamics and the strength of selection for resistance.

The first key implication of our work is that as well as suppressing growth and colonization by invading strains [23, 52], the gastrointestinal microbiota can inhibit antibiotic resistance evolution. There are several possible ecological mechanisms by which the microbiota may suppress growth of invading lineages [53], such as niche and nutrient competition [54], direct killing via bacteriocins [22, 55], phage production [56] or by changing the concentration of compounds such as primary and secondary bile acids [57, 58]. It was not our aim to pull apart the mechanisms by which resident microbiota suppress invading bacteria (studied in more detail elsewhere; [55, 59]). Nevertheless, our data on community structure indicate resident *Enterobacteriaceae* including *E. coli* had a competitive advantage in our system, potentially explaining suppression of the focal strain. This was further evidenced by the advantage of transconjugants carrying plasmids from resident *E. coli* (S5 Fig) and resident *E. coli* over our ancestral focal strain in competition experiments (S7 Fig). The competitive advantage of resident *E. coli* extended to ampicillin-free conditions in pure culture. Possible contributors to this include a type VI secretion system with various effectors we detected in the genomes of the human donor 1 and 3 *E. coli* isolates, and the a colicin plasmid present in human donor 1 *E. coli* isolates (Fig 4). In supernatant experiments, we did not find evidence of direct inhibition via phages in the community samples (see S1 Methods). Crucially, no matter how interactions with the microbiota suppress growth of an invading lineage, we expect the reduced population size, growth and replication to in turn reduce the supply of new genetic variation. This is consistent with *in vitro* work with malaria parasites showing competition between two species under resource limitation impeded drug resistance evolution [60] and previous studies with *Pseudomonas fluorescens* showing that a eukaryotic predator [61] or a bacteriophage [62] can suppress the emergence of antibiotic resistance. Thus, suppression of an invading lineage via interactions with resident microbiota may frequently have a knock-on effect on resistance evolution.

A second key implication is that resident microbiota modified selection on antibiotic resistance in our focal strain. The stronger effect of ampicillin on focal strain growth in the presence of resident microbiota (Fig 1; S1 Table) indicates resistance would have been more beneficial here. In support, the benefits of resistance plasmid acquisition were greatest in the presence of resident microbiota (S5 Fig). This is counter to the expectation that antibiotics may be less effective in more dense communities due to an ‘inoculum effect’ [63]. We found inhibition of our focal strain was indeed altered by very high *E. coli* abundance in pure cultures (S8 Fig), although there was still considerable inhibition even at the highest densities. This indicates such inoculum effects were weaker in the presence of species-rich communities than in pure cultures of *E. coli*, and/or were counterbalanced by opposing effects of community-level interactions on ampicillin inhibition of *E. coli*. By contrast, chromosomal resistance mutations that emerged in the absence of resident microbiota (and which conferred relatively small increases in resistance in pure culture) were no longer beneficial in the presence of resident microbiota, indicating a larger change in resistance is needed to overcome the relatively strong effect of ampicillin here. This complements recent work showing natural communities from pig faeces can increase costs of antibiotic resistance for individual species [12], and that costs of CRISPR-based phage resistance can be amplified by interactions with other bacterial species [64]. More generally, this supports the notion that community-level interactions modulate the costs and benefits of antibiotic resistance via mechanisms that are only just beginning to be understood [65].

A third key implication of our data concerns the genetic and environmental constraints on horizontal gene transfer that determine whether or not the growth-suppressive effects of the microbiota are counter-balanced by horizontal transfer of beneficial genes. The unavailability of known resistance genes to the invading focal strain in the community from human donor 3 was because they were integrated in the chromosome. The plasmid we isolated from the human donor 1 community was conjugative, but transfer depended on the abiotic conditions. This suggests the potential for plasmid transfer to allow invading lineages to overcome the suppressive effects of the microbiota depends critically on whether they are conjugative (which can be predicted from sequence data) and on the sensitivity of conjugative transfer to local physical conditions (which is harder to predict from sequence data). Consistent with this, previous research has shown conjugative transfer in *E. coli* and other species is sensitive to the physical experimental conditions [51, 66]. Furthermore, mating pair formation machinery, usually encoded by the plasmid, in some cases promotes biofilm formation, which can in turn promote the spread of plasmids [67]. This raises the question of whether some plasmids have evolved to manipulate the physical structure of bacterial populations to promote transfer. Despite these constraints, plasmids are clearly sometimes transferred *in vivo*, as has been observed in animal models [68, 69] and human gut microbiomes [33,44,45]. In line with plasmids being key vectors of beta-lactamases [70], the conjugative plasmid we identified was highly effective in terms of resistance. This suggests plasmid-borne resistance will be under strong positive selection once established and can spread rapidly via clonal expansion. However, our experiment showed the initial horizontal transfer required for such spread is sensitive to genetic and abiotic constraints.

Our approach allowed us to isolate the effect of interactions between diverse microbial communities and a focal *E. coli* strain. Our amplicon sequence data showed the communities had a representative taxonomic composition for healthy human donors. There were still diverse communities present at the end of the seven-day experiment, albeit with a change in the relative abundance of different taxa. The observed rise of *Enterobacteriaceae* and *Bacteroidaceae* has been seen in other experiments with gastrointestinal communities (e.g. [71]) and might be explained by the nutrient content of the medium, micromolar oxygen levels or such *in vitro* systems favouring faster population growth [72]. In conditions where antibiotics are applied at higher concentrations or affect a greater fraction of extant taxa, we can expect stronger shifts in community composition in response to antibiotic treatment. We used a sub-lethal concentration in our main experiment to allow us to track invasion and growth by the focal strain. Nevertheless, in competition experiments at higher antibiotic concentrations we saw similar outcomes in that plasmids from the resident microbiota were highly beneficial whereas chromosomal mutations were not (S5 Fig). More importantly, the shift in community composition over the seven-day experiment does not explain the observed suppression of the focal strain, because this was already visible after one day.

Although our experimental system likely differs from the gastrointestinal tracts these bacteria were isolated from in ways that affect community composition, cultivating them *in vitro* allowed us to quantify the effect of species-rich communities sampled from gastrointestinal tracts on resistance evolution of a relevant opportunistic human pathogen. A key limitation of our study is the sample size (three human donors, one focal strain, one antibiotic). Some outcomes might change with different types of resident microbiota or different types of plasmids (explored in the Supplementary Model). Nevertheless, we observed a qualitatively consistent suppression of the focal strain across the three human donors, which was always stronger in the presence of ampicillin, and in some cases was associated with colonization resistance (extinction of the focal strain). Additionally, we chose *E. coli* and ampicillin because they are both important for understanding resistance evolution in nature and share some important properties in this respect with other bacteria and antibiotics (our rationale is explained further in the introduction). Despite the low sample size, we observed a qualitatively consistent suppression of the focal strain across the three human donors, which was always stronger in the presence of ampicillin, and in some cases was associated with colonization resistance (extinction of the focal strain). A key challenge for future work will be to determine how such effects are modified *in vivo* by local spatial structure [73] and immune responses [74]. Indeed, interactions mediated via the host immune system are another possible mechanism of colonization resistance [75–77].

In conclusion, we showed species-rich microbial communities sampled from human gastrointestinal tracts can suppress growth and resistance evolution of an invading lineage. Given the variety and likely common occurrence of mechanisms that can generate such suppression of invaders (e.g. resource competition), these types of effects are probably common in species-rich communities such as the mammalian gastrointestinal tract. Crucially, resident microbiota also altered the strength of selection for resistance (ampicillin was more suppressive for the focal strain in community treatments) and the fitness effects of individual genetic changes (high-level resistance plasmids became more beneficial in the community treatments, but low-level resistance mutations became less beneficial). Our other data and simulations showed whether or not the growth-suppressive effects of resident microbiota are counter-balanced by beneficial horizontal gene transfer depends on genetic and environmental constraints that can impede the spread of resistance plasmids. This has important implications for the prediction of resistance evolution from genetic and metagenomic data, such as those widely collected through surveillance efforts [78, 79]: identifying mobile resistance genes in a diverse community is not enough to predict resistance evolution, requiring in addition information about genetic and environmental constraints on *in situ* transfer dynamics.

## Material and Methods

### Bacterial strains

We used *Escherichia coli* K12 MG1655 carrying a Streptomycin resistance mutation (*rpsL* K43R) as the focal strain. Two days prior to the experiment we streaked the focal strain on LB-agar (Sigma-Aldrich, Buchs, Switzerland) and incubated overnight at 37 °C. To incubate the focal strain cultures anaerobically prior to the microcosm experiment, we prepared 42 Hungate tubes (VWR, Schlieren, Switzerland) with LB (Sigma-Aldrich), which was supplemented with 0.5 g/l L-Cysteine and 0.001 g/l Resazurin (reducing agent and anaerobic indicator respectively), flushed the headspace with nitrogen, sealed the tubes with a rubber stopper and autoclaved them. One day before the experiment, we randomly picked 42 colonies and inoculated them in the 42 Hungate tubes containing anaerobic LB, and incubated at 37 °C overnight with 220 rpm shaking. We then used these 42 independent cultures of the focal strain to inoculate the main experiment described below.

### Human microbiome samples

We used three stool samples from anonymous, consenting human donors. All samples were collected at the Department of Environmental Systems Science, ETH Zürich on the 15^th^ of May 2018. Inclusion criteria were: older than 18 years, not obese, not recovering from surgery and no antibiotics in the last six months. The sampling protocol was approved by the ETHZ Ethics Commission (EK 2016-N-55). Each sample was collected in a 500 ml plastic specimen container (Sigma-Aldrich) and kept anaerobic using one AnaeroGen anaerobic sachet (Thermo Scientific, Basel, Switzerland). The three samples used for the experiment were randomly selected from a larger number of donated samples. We collected the samples in the morning before the experiment and kept them for maximum one hour before processing. To prepare faecal slurry from each sample, we re-suspended 20 g of sample in 200 ml anaerobic peptone wash (1 g/l peptone, 0.5 g/l L-Cysteine, 0.5 g/l bile salts and 0.001 g/l Resazurin; Sigma-Aldrich) to prepare a 10% (w/v) faecal slurry. We then stirred the slurry for 15 min on a magnetic stirrer to homogenize, followed by 10 min of resting to sediment. At this point we removed 100 ml of each faecal slurry (’fresh slurry’), which we used later to re-introduce the resident microbial community to sterilized slurry (for the community treatments). To sterilize the faecal slurries, we transferred 100 ml to a 250 ml Schott bottle, flushed the headspace with Nitrogen gas, sealed them with rubber stoppers and autoclaved for 20 min at 121 °C.

### Inoculating anaerobic gut microcosms, sampling and bacterial enumeration

For the start of the experiment (S1 Fig), we filled 42 Hungate tubes with 7 ml of basal medium, which was based on earlier studies [80, 81] with some modifications (2 g/l Peptone, 2 g/l Tryptone, 2 g/l Yeast extract, 0.1 g/l NaCl, 0.04g K_2_HPO_4_, 0.04 g/l KH_2_PO_4_, 0.01 g/l MgSO_4_x7H_2_O, 0.01 g/l CaCl_2_x6H_2_O, 2g/l NaHCO_3_, 2 ml Tween 80, 0.005 g/l Hemin, 0.5 g/l L-Cysteine, 0.5 g/l bile salts, 2g/l Starch, 1.5 g/l casein, 0.001g/l Resazurin, pH adjusted to 7, addition of 0.001g/l Menadion after autoclaving; Sigma-Aldrich), and for the subsequent re-inoculation cycles with 6.5 ml of basal medium. We flushed the head space of each tube with nitrogen gas, sealed it with a rubber septum and autoclaved to produce anaerobic microcosms containing only basal medium.

On day one of the experiment, we introduced faecal slurry and antibiotics to each tube according to a fully factorial design (S1 Fig), with three replicate microcosms in each combination of Human Donor (1, 2 or 3), Community (present or absent) and Antibiotic (with or without). In the community-free treatments, we added 850 µl of sterile slurry. In the community treatments, we added 350 µl of fresh slurry and 500 µl of sterilized slurry. In the antibiotic treatment, we added ampicillin to a final concentration of 7.2 µg/ml, approximating the IC90 (concentration required to reduce growth by 90%) for the focal strain; this was introduced 2h after the focal strain had been inoculated (8 µl of focal *E. coli* from one of the 42 overnight cultures introduced at 0h; approximately 1:1000 dilution). As a control treatment testing for the effect of sterilized slurry, we also inoculated the focal strain into three replicate microcosms containing only the basal medium (supplemented with 850 µl of peptone wash to equalize the volume with the slurry treatments), and we did this with and without antibiotic treatment. We incubated all microcosms at 37 °C in a static incubator. After 24h we transferred a sample of 800 µl from each microcosm to a new microcosm (containing basal medium plus 500 µl of sterile slurry from the corresponding human donor in the community and community-free treatments, or basal medium plus peptone wash for the basal medium treatment, supplemented with ampicillin at each transfer in the antibiotic treatments), and we repeated this for seven days.

To estimate the abundance of the focal strain during the experiment, we used a combination of selective plating and colony PCR. For selective plating, we serially diluted the samples and plated them on Chromatic MH agar (Liofilchem, Roseto degli Abruzzi, Italy), which allowed us in a first step to discriminate *E. coli* from other species based on colony colour. By supplementing these agar plates with Streptomycin (200 µg/ml), to which our focal strain is resistant, we selected against other *E. coli* that were not resistant to Streptomycin. To screen for variants of our focal strain that acquired resistance to Ampicillin during the experiment, we additionally plated each sample onto the same agar supplemented with both Streptomycin (200 µg/ml, Sigma-Aldrich) and Ampicillin (8 µg/ml, Sigma-Aldrich). We did this after every growth cycle. Despite initial screening of microbiome samples revealing no resident *E. coli* that could grow on our selective plates, later in the experiment we found such bacteria to be present in some samples (that is, non-focal strain *E. coli* that could grow on our plates and were presumably very rare at the beginning of the experiment). To discriminate between these *E. coli* and our focal strain, we used colony PCR with specific primers (forward (5’-AGA CGA CCA ATA GCC GCT TT-3’); reverse (5’-TTG ATG TTC CGC TGA CGT CT-3’)). For colony PCR, we picked 10 colonies randomly for each time point and treatment. The PCR reaction mix consisted of 2x GoTaq green master mix, 2.5 μM of each primer and nuclease free water. The thermal cycle program ran on a labcycler (Sensoquest, Göttingen, Germany) with 6 min 95°C initial denaturation and the 30 cycles of 95°C for 1 min, 58°C for 30s, 72°C for 35s and a final elongation step of 72°C for 5 min. For gel electrophoresis we transferred 5 μl of the PCR reaction to a 1.5% agarose gel stained with SYBR Safe (Invitrogen, Thermo F. Scientific) and visualized by UV illumination. Focal strain abundance was then estimated by multiplying the frequency of the focal strain determined by colony PCR with the total colony count for each plate. To account for the possibility that the focal strain was still abundant in populations where we found 0/10 colonies (that is, where it could have been rare relative to resident *E. coli* but still present), we additionally screened the DNA extracted from the community (described below) of the final time point by PCR (as described above for the colony PCR, but using 30 ng of DNA as template). We did this for all microcosms from the community treatment and detected PCR products in all cases where we detected focal strain by plating and colony PCR, and none of the cases where we did not, consistent with our analysis of individual colonies and suggesting that in those microcosms the focal strain had been completely excluded during the experiment.

To estimate total microbial abundance in each microcosm supplemented with the microbiome we used flow cytometry. We diluted samples by 1:10000 with phosphate buffered saline (PBS) and stained them with SYBR Green (Invitrogen, Thermo F. Scientific). We used a Novocyte 2000R (ACEA Biosciences Inc., San Diego, CA, USA), equipped with a 488 nm laser and the standard filter setup for the flow cytometric measurements.

We froze samples after every transfer from each microcosm at −80 °C, and at the end of the experiment we isolated two sets of focal strain colony isolates for sequencing. First, we randomly picked a single focal strain colony isolate from each microcosm where the focal strain was detected at the end of the experiment (from Streptomycin plates; *n*=33). Second, we randomly picked a single ampicillin-resistant colony isolate of the focal strain from each of the 8 populations at the end of the experiment where they were detected (from Streptomycin + Ampicillin plates; *n*=8). We grew each colony isolate overnight in LB (with Ampicillin for the ampicillin-resistant isolates), mixed 1:1 with 50% glycerol and stored at −80 °C.

### Whole genome sequencing and Bioinformatics

We sequenced all of the randomly selected (*n*=33) and ampicillin-resistant (*n*=8) focal strain colony isolates (S7 Table). Prior to DNA extraction, we grew each isolate overnight in LB, then concentrated the liquid cultures by centrifugation and extracted DNA with the Magattract kit (Qiagen, Hilden;Germany) according to the manufacturer’s protocol. Quality and quantity of DNA was assessed with Nanodrop (Thermo Fisher) and Qubit (Thermo Fisher). At the Functional Genomics Center (ETH Zürich / University of Zürich) DNA was processed with Nextera library prep kit and sequenced on the Illumina Hiseq 4000 platform. We filtered raw sequencing reads with trimmomatic version 0.38 [82] and mapped the reads to the reference with snippy version 0.9.0 (https://github.com/tseemann/snippy) to detect variants. Deletions were identified by using the coverage analysis tool of CLC Genomics Workbench 11 (Qiagen) with a read threshold value of 0 and a window size of 1. We identified IS elements with ISfinder web server (database from July 2018) (Siguier 2006) in the ancestor genome and used these sequences to detect IS movements in the evolved strains with ISMapper version 2.0 [84].

We additionally sequenced 12 ampicillin-resistant resident *E. coli* colony isolates (not the focal strain, S7 Table). We isolated these colony isolates from microcosms filled with faecal slurry from human donors 1 and 3 at the final time point of the experiment (6 for each donor, each from a different microcosm). We picked, grew and sequenced these colony isolates as described above for focal strain isolates. We then made de Novo assemblies of the resulting sequences with spades version 3.13.0 [85] and annotated them with prokka version 1.13.7 [86].

Additionally, we sequenced one of these resident, resistant *E. coli* isolates from both human donors (1 & 3) with the Oxford nanopore long-read sequencing platform MinION at University Hospital Basel, Switzerland. These genomes were assembled using Unicycler v0.4.8 with a hybrid assembly approach combining MinION and Illumina reads. Assembly statistics can be found in S6 Table. We screened for known antibiotic resistance genes and the presence of plasmid replicons by a local blast query against the resfinder (downloaded Oct. 2018) [87] and plasmidfinder (downloaded Oct. 2018) [47], which is a repository of whole-plasmid sequences from members of the *Enterobacteriaceae*. To identify genes that are involved in mating pair formation or mobilization of the plasmid we screened the genome annotation files and blasted potential candidate genes against the NCBI nucleotide database to verify them.

### Mating experiments with plasmids from resident *E. coli*

We aimed to determine whether the resident *E. coli* colony isolates could act as plasmid donors for our focal strain (transferring antibiotic resistance plasmids via conjugation). Because the replicate *E. coli* colony isolates that we sequenced from each human donor (1 and 3) were almost identical on the DNA sequence level (S2 Table), we randomly chose one colony isolate from each human donor as a potential plasmid donor strain. We used a focal strain as the potential plasmid recipient in these experiments, which was only different from the focal strain used in the main experiment by addition of a dTomato marker and a Chloramphenicol resistance cassette (enabling us to detect transconjugants by selective plating). In the first set of mating experiments, we grew overnight cultures of the potential plasmid donor strains and the potential plasmid recipient strain in LB at 37 °C with shaking. We then pelleted the overnight cultures by centrifugation (3000 rpm, 5 min), washed them twice with PBS and resuspended them in 300 µl PBS. We then mixed the donor and recipient strains 1:1 (v:v), and transferred 100 µl of this mixture to each of three replicate LB plates, and incubated the plates for 6h at 37 °C. After that, we washed the cells off the plate with 500 µl PBS and streaked out 50 µl of this plate wash on LB plates supplemented with Ampicillin and Chloramphenicol. To verify plasmid uptake by the transconjugants, we used a colony-PCR screen with primers targeting the recipient strain (same primer set we used above to identify the focal strain) and three additional primer sets targeting the beta-lactamase gene *bla_Tem_-1b* (blaFW: 5’-TGCAACTTTATCCGCCTCCA-3’; blaRV 5’-TTGAGAGTTTTCGCCCCGAA-3’), the *traC* gene (traFW 5’-TCGATAAACGCCAGCTGGTT-3’; traRV 5’-AGGTGAAAACCCACAGCGAA-3’) and the replicon *IncFIC (FII)* (IncFW 5’-CACCATCCTGCACTTACAATGC-3’; IncRV 5’-TCAGGCCCGGTTAAAAGACA-3’) with the same PCR reaction mix and settings as described above.

To test whether environmental conditions affected conjugation efficiency, we performed a second set of mating experiments in four different conditions: solid LB agar, liquid LB, liquid anaerobic LB and anaerobic sterile faecal slurry. We prepared the liquid LB (Sigma-Aldrich) and LB agar (Sigma-Aldrich) for all treatments in the same way (independent of whether they were aerobic or anaerobic), supplementing them with 0.5 g/l L-Cysteine and 0.001 g/l Resazurin. For the anaerobic LB treatment, we transferred 0.9 ml LB to each Hungate tube, flushed the headspace with nitrogen and sealed it with a rubber stopper before autoclaving. For the anaerobic sterile faecal slurry treatment, we added 0.45 ml LB to each Hungate tube and 0.45ml thawed slurry under anaerobic conditions, before flushing the headspace with Nitrogen, sealing and autoclaving. Prior to each mating assay, recipient and donor strains were inoculated, washed, concentrated and mixed exactly the same way as described above for the first set of mating experiments. For each solid and liquid treatment four replicates were inoculated with 100 μl of the 1:1 donor recipient mix, either under aerobic or anaerobic conditions according to the respective treatment and all tubes and plates were incubated for 6h under static conditions at 37 °C. We stopped the mating assay by either vortexing the liquid cultures or washing off the cells from the plates with 1 ml of PBS. 100 μl of each bacterial suspension was then plated on selective agar plates containing either Chloramphenicol to count total number of recipient cells or a mix of Chloramphenicol and Ampicillin to count the number of transconjugants. After 24h we calculated transconjugant frequencies by dividing colony-forming unit (C.F.U.) counts of plasmid-positive colonies by the total C.F.U. count of recipient cells [69].

We measured susceptibility to ampicillin for the ancestral focal strain from the main experiment, the focal strain used in the conjugation experiment (with the dTomato tag), one focal strain transconjugant (with the plasmid from the resident *E. coli* isolate of human donor 1), the resident *E. coli* isolate of human donor 1 (carrying the same plasmid), and all eight ampicillin resistant focal strain isolates from the main experiment. We did this by measuring OD600 after 24h incubation at various concentrations of ampicillin. We prepared overnight cultures of each isolate in a randomly organized master plate, and then inoculated the susceptibility assay using a pin replicator to transfer approximately 1 μl of the overnight cultures to assay plates filled with 100 μl of 0-60 μg/ml ampicillin per well. We measured OD at 0h and after 24h with a NanoQuant infinite M200Pro platereader (Tecan).

### Amplicon sequencing

We thawed samples of fresh faecal slurry from 0h and samples from each microcosm in the community treatments after 24h and 168h on ice and homogenised them by vortexing. We concentrated each slurry sample by centrifuging 1.5 ml of each sample at 3000 rpm directly in the bead beating extraction tube, before removing the supernatant and repeating this step, resulting in a total volume of 3 ml of each slurry sample. We then extracted the DNA from this concentrate following the protocol of the powerlyzer power soil kit (Qiagen). DNA yield and quality were checked by Qubit and Nanodrop.

We amplified the V3 and V4 region of the 16S rRNA gene with three slightly modified universal primers [88] with an increment of a 1-nt frame shift in each primer [89] to increase MiSeq sequencing output performance between target region and Illumina adapter. The target region was amplified by limited 17 cycle PCR with all three primer sets in one reaction for each sample. We cleaned up PCR products and in a second PCR adapters with the Illumina barcodes of the Nextera XT index Kit v2 were attached. We checked 10 randomly selected samples on the Tapestation (Agilent, Basel, Switzerland) for the proper fragment size. We quantified library by qPCR with the KAPA library quantification Kit (KAPA Biosystems, Wilmington, MA, USA) on the LightCycler 480 (Roche, Basel, Switzerland). We normalized quantified samples, pooled them in equimolar amounts and loaded the library with 10% PhiX on the Illumnina MiSeq platform with the Reagent Kit V3 at the Genetic Diversity Center (ETHZ).

Sequencing reads were trimmed on both ends with seqtk version 1.3 (https://github.com/lh3/seqtk) and amplicons were merged into pairs with flash version 1.2.11 [90]. USEARCH version 11.0.667 [91] was then used to trim primer sites and amplicons were quality filtered using prinseq version 0.20.4 [92]. We clustered the quality filtered sequences into zero-radius operational taxonomic units (ZOTU) using USEARCH. We used Sintax implemented in the USEARCH pipeline with the SILVA database [93] for the taxonomic assignment of the ZOTU sequences. For the analysis of taxonomic data including plotting Shannnon diversity, relative proportions of taxa and to generate the PCoA plots, we used the Phyloseq package in R [94]. To estimate the frequency of *E. coli* relative to total *Enterobacteriaceae*, we first isolated all ZOTUs assigned to *Enterobacteriaceae* and divided these reads into *Escherichia coli* and other bacteria. We blasted both groups against the SILVA database to check if the reads were assigned to the right group.

### Statistical analyses

We used R 3.5.1 [95] for all analyses. To test whether focal strain abundance differed between treatments, we used a generalized linear mixed effects model with the glmmadmb function of the glmmADMB package, with zero inflation and a Poisson error distribution [96]. For this analysis we excluded the basal medium treatment and used time, antibiotic (with / without), resident microbial community (with / without) and human donor (1 / 2 / 3) as fixed effects and replicate population as a random effect. The model was reduced by removing non-significant (P>0.05) interactions using F-tests. P values for interaction terms in the reduced model were obtained with type II Wald chisquare tests. To test whether there was an inhibitory effect of sterilized slurry on the focal strain, we used the glmer function of the lme4 [97] R package. For this analysis we included only community-free and antibiotic-free treatments, with focal strain abundance as the response variable, donor as a fixed effect, replicate population as a random effect, and a Poisson error distribution. After finding interactions between the effects of resident microbiota depending on human donor and antibiotic, we analyzed subsets of the dataset to look at ampicillin-free treatments only and individual human donors, using the same approach as for the main model.

To analyse the effects of ampicillin and community presence/absence on the competitive fitness of mutants and transconjugants (see S1 Methods), we used a linear model with the lmp function of the lmperm package [98]. Here we took relative fitness for the respective mutant or transconjugant as the response variable and antibiotic concentration and community presence/absence as fixed effects, testing factor effects by permutation test (accounting for the non-normal distribution of our fitness data).

To analyse differences in total bacterial abundance in the community treatments, we used a linear mixed effects model with the lmer function of lme4 [97], with a Poisson error distribution. Time, donor and antibiotic were fixed effects and replicate population a random affect. To analyse variation of Shannon diversity, we used a linear mixed effects model with the lme function of nlme [99]. We excluded time point 0h from the analysis, and included Time, Donor and Antibiotic as fixed effects and replicate population as a random effect. To analyze similarities of microbiome samples based on the 16S rRNA data, we applied the Bray-Curtis distance metric with the ordinate function of the Phyloseq R package to get coordinates for the principal coordinate analysis (PCoA). On this dataset we ran a permutational multivariate analysis of variance (PERMANOVA) with the adonis function of vegan [100], using the distance matrix obtained from the PCoA analysis but omitting time point 0h. We did this separately for time points 24h and 168h.

## Supporting information

S1 Model

## Acknowledgements

We thank the Genetic Diversity Center (ETH Zurich), the Functional Genomics Center (ETH Zurich / University of Zurich), Jean-Claude Walser for sequencing and bioinformatics support, Adrian Egli and Helena Seth-Smith from University hospital Basel for help with MinION sequencing, Jana Huisman for help with simulations and Vera Beusch for help compiling plasmid transfer rate estimates from literature.

## Supplementary Figure legends

**S1 Fig. Summary of experimental evolution in faecal slurry**. Treatments consisted of basal medium only, basal medium supplemented with sterilized faecal slurry (without the resident microbial community) from one of three human donors, or basal medium supplemented with sterilized faecal slurry to which the resident microbial community had been reintroduced (with community). After inoculation, all treatments were incubated for 2h at 37 °C, before 7 μg/ml ampicillin was added in the antibiotic treatment. Every 24h, we sampled each microcosm and transferred an aliquot to fresh medium (either basal medium or sterilized faecal slurry) with or without antibiotics. We serially diluted each sample and spread it on chromatic agar plates with or without antibiotics to quantify focal strain abundance (verified by colony PCR) and to screen for resistance. We sequenced focal strain isolates from the final time point and investigated community composition by 16S rRNA gene amplicon sequencing. We monitored total bacterial abundance in the community treatments by flow cytometry.

**S2 Fig. Variable similarity of microbial communities across time and treatment groups**. Each panel shows samples from a single human donor, with the same axes used in each panel. Points show the initial sample (0h) and microcosms from 24h and 168h with and without antibiotics (legend at right). Similarities between communities were calculated by Bray-Curtis distance and plotted using Principal Coordinate Analysis (see Methods).

**S3 Fig. Abundance of sequences associated with the focal strain, total *E. coli* and the resistance plasmid from the microbiota of human donor 1, inferred with qPCR**. Each panel shows the copy number of sequences detected with primers specific for the focal strain, total *E. coli* and the resistance plasmid (see legend; further details of primers in S1 Methods) at timepoint 0h (left panel), 24h (middle panel) and 168h (right panel). Each point shows the mean of three technical replicates. Reactions where no amplification was detected are shown at 10^0^. We expect plasmid copy number to reflect the abundance of plasmid donor cells, because coverage analysis of whole genome sequencing data indicated a copy number per cell of ∼1. For the focal strain and total *E. coli*, the copy number of sequences does not necessarily reflect the total number of cells of each type, but changes in strain abundance over time would nevertheless be expected to result in strongly correlated changes in sequence copy numbers over time.

**S4 Fig. Agarose gel electrophoresis picture of the PCR products specific for plasmid genes and a chromosomal marker of the focal strain.** We used these primer sets to verify plasmid uptake of the transconjugants. Primers are given in the main text in the methods section.

**S5 Fig. Competitive fitness of transconjugants and mutants relative to the ancestral focal strain in the presence and absence of resident microbial communities with no, sub-MIC and supra-MIC ampicillin**. (A) Final cell densities of competing strains (see legend; Transconjugant is a transconjugant of the focal strain carrying the plasmid from Human Donor 1, in the left panel; Mutant is an evolved isolate with increased ampicillin-resistance from the community-free treatments with slurry from Human Donors 1, in the middle panel or Human Donor 3, in the right panel; Ancestor is the respective ancestral focal strain). Data are shown after 24h of competition in sterile slurry or community treatments, with and without low or high concentrations of ampicillin (*x*-axis). (B) Fitness of the transconjugant or mutant relative to the ancestor, calculated as the difference of their Matlhusian growth parameter in the same experiment. In both panels the three points show three replicates of the experiment.

**S6 Fig. Competitive fitness of transconjugants (carrying the plasmid from resident *E. coli* of Human Donor 1) relative to evolved isolates (from community-free treatments with faecal slurry from Human Donor 1, left, and Human Donor 3, right).** In each panel relative fitness of the transconjugant strain is shown as the difference in malthusian growth parameters compared to the respective evolved isolate (see S1 Methods). Competitions were done in sterile faecal slurry or the presence of the resident microbial community, and with no, low or high ampicillin concentrations (*x*-axis). Each point shows a different replicate.

**S7 Fig. Abundance of the focal *E. coli* strain and resident *E. coli* strains isolated from Human Donors 1 and 3 (see legend) in monoculture (left) and in co-culture (right).** Each strain was grown in monoculture in the absence of antibiotics and each co-culture combination was grown in the presence and absence of ampicillin (*x*-axis). Each point shows a different replicate.

**S8 Fig. Effect of starting bacterial density on growth inhibition by ampicillin (’inoculum effect’):** Changes in bacterial abundance over 24h are shown using three different quantification methods (optical density, top panel; plating and C.F.U. counting, middle panel; flow cytometry, bottom panel). In each panel the change in abundance is shown for four starting densities (see legend) and at four antibiotic concentrations. In each panel the change between 0h and 24h is shown (in OD in the top panel, in C.F.U./ml in the middle panel, and in recorded events/ml in the bottom panel). Each point shows the mean of three replicates, error bars show 1 s.d.

