## Supplementary material for "Resident microbial communities inhibit growth and antibiotic resistance evolution of *Escherichia coli* in human gut microbiome samples": S1 Methods

### Supporting material and methods

*Quantitative PCR analysis of* E. coli *abundance*

To estimate changes in the abundance of genetic sequences associated with the focal strain, total *E. coli* and the resident resistance plasmid over time in community treatments, we used qPCR. We used a specific primer for our focal strain (fw primer: 5'-GCGGTAATCCGGAAAACTCA-3'; rv primer: 5'-GTACCCGGTATCGGTGCTTC-3'), a primer for all *E. coli* targeting the *ybbW* gene [1] (fw primer: 5'-TGATTGGCAAAATCTGGCCG-3'; rv primer: 5'-GAAATCGCCCAAATCGCCAT-3') and a primer pair specific for the plasmid found in human donor 1 (targeting parts of the beta-lactamase gene and the adjacent intergenic region, fw primer; 5'-CGCTGAGATAGGTGCCTCAC-3'; rv primer: 5'-TAGCGTAGGATTCGGGGTCT-3'). We performed each qPCR reaction in a final volume of 20 µl with 10 µl of Takyon No Rox SYBR mastermix dTTP blue (Eurogentec, Seraing, Belgium), 100 nM of each primer and 2.5 µl of DNA template from extraction for amplicon sequencing as described in the material and methods section. We ran the qPCR on a LightCycler 480 (Roche, Basel, Switzerland) with the standard protocol including one activation cycle of 95°C for 3 min, followed by 40 cycles at 95°C for 10 s, 60°C for 20 s and 72°C for 20 s. Standard curves were run in duplicates and all samples in triplicates. We included a negative control for each primer pair using 5 µl of water. We generated the standard curves for the primer sets of the focal strain and total *E. coli* abundance using a 10-fold dilution series of DNA extracted from *E. coli* K12 MG1655 covering a range from 50 to $5\times10^7$ genome equivalents, calculated based on genome size. For the plasmid standard curve, we first extracted the plasmid DNA from the transconjugant with the ZR BAC DNA extraction kit (Zymo, Irvine, California, USA) following the manufacturer's protocol and then prepared a 10-fold dilution series ranging from 50 to $5\times10^7$ plasmid equivalents, calculated based on plasmid size. We exported crossing point (Cp) values calculated by the LightCycler 480 II software V1.5 and used the Cp values for each standard to plot a standard curve by assigning to each Cp value a concentration. The standard curve for the focal strain primers was (y=-3.48+43.85, $R^2$=0.99), for the *E. coli* primer set (y=-3.53x+44.14, $R^2$=0.99) and for the plasmid primer set (y=-3.49+44.47, $R^2$=0.99).

*Competition assays with mutants and transconjugants in presence of the community*

To test whether resistance mutations from community-free treatments and plasmids from resident microbiota provided fitness benefits to the focal strain in the presence of communities, we used competition experiments. In each experiment we cultivated the ancestral focal strain together with an evolved ampicillin-resistant mutant (one representative mutant from human donor 1, community-free-with-ampicillin treatment, replicate 1, and one from human donor 3, community-free-with-ampicillin treatment, replicate 1) or we cultivated the ancestral focal strain used to isolate transconjugants together with the transconjugant obtained from the mating assay. We did this anaerobically as in the main experiment, with and without the community and at three different ampicillin concentrations (zero, the same as in the main experiment, and 3× the IC90 of the focal strain). We pre-cultivated independent cultures of each strain as in the main experiment and prepared faecal slurry treatments as described in the in the material and methods section, but using frozen slurry. We inoculated competitions with 8 µl of pre-culture of each strain, and added zero, 7.2 µg/ml or 21,6 µg/ml ampicillin before incubation for 24h at 37°C. We estimated initial and final cell densities by plating on selective agar with streptomycin (for the ancestral focal strain and evolved ampicillin-resistant mutants) or chloramphenicol (for the ancestral focal strain used to make transconjugants and for transconjugants), before using colony PCR as above to estimate the frequency of each strain. We then estimated the Malthusian growth parameter for each strain to estimate the competitive fitness of mutants and transconjugants relative to their respective ancestors [2].

*Competition assays with focal strain and resident* E. coli *in absence of community*

To test whether resident *E. coli* strains had a competitive advantage against the focal strain in the absence of a species-rich community, we performed competition assays in the presence and absence of ampicillin. We cultivated the focal strain (a variant with a chloramphenicol and dTomato selection marker) either as monoculture or in co-culture with resident isolates (one isolate from human donor 1, with-community-with-ampicillin treatment, replicate 1, and one isolate from human donor 3, with-community-with-ampicillin treatment, replicate 1). We pre-cultured and inoculated each culture as in the main experiment in basal-medium-only conditions, adding 8 µl of pre-culture for monoculture treatments and 8 µl of each strain for co-cultures. We estimated initial (0h) and final (24h) population densities by plating on selective agar with chloramphenicol (for the focal strain) or ampicillin (for resident *E. coli* strains), before estimating relative fitness as above.

*Test for inoculum-effect*

To investigate the effect of starting density on inhibition by ampicillin, we grow bacteria from four starting densities ($10^9$, $10^8$, $10^7$, $10^6$ C.F.U./ml) at four ampicillin concentrations (0, 4, 8, 16 µg/ml). We grew independent cultures of *E. coli* MG1655 in 200 µl of LB in a 96-well microplate overnight at 37°C in a shaking incubator. We then diluted cultures by varying degrees into 200 µl of LB, in triplicate at each antibiotic concentration. For the highest density ($10^9$), we did not dilute the overnight cultures but instead centrifuged them (10 min at 3000rpm) and resuspended the pellet in 200 µl of LB medium with each antibiotic concentration, before transfer to the same assay microplate as the other starting density treatments. We incubated the plate at 37°C in the shaking incubator, measuring OD at 0h and 24h with a NanoQuant infinite M200Pro platereader (Tecan, Männedorf, Switzerland). We used the same experimental set-up (but a different temporal block), to quantify the inoculum effect using alternative methods of counting bacteria: plating (C.F.U. count) and flow cytometry. For plating, we diluted the different treatments in PBS and plated them on LB agar in different dilutions according to the density of the sample (dilutions between $10^0$ and $10^7$) at 0h and 24h after incubation. Flow cytometry samples were also diluted in PBS (dilutions between $10^0$ and $10^6$), stained with SYBR green (Invitrogen, Thermo F. Scientific) and measured with a Novocyte 2000R flow cytometer (ACEA Biosciences Inc., San Diego, CA, USA).

*Supernatant assay for potential phage killing*

We also screened the initial faecal slurries from each human donor for evidence of lysis of our focal strain, which might be caused by lytic phages. We centrifuged each faecal slurry (10 min, 2000 g), filtered the supernatant with cellulose (Melitta, Egerkingen, Switzerland), then 0.45 and 0.22 μm syringe filters, then supplemented with $MgSO_4$ and $CaCl_2$ (final concentration 8 μM), and stored at 4 °C. We then made lawns of the focal strain on agar and added 10 μl of supernatant extract from each human donor and incubated 24h at 37 °C. We never observed any plaques or evidence of lysis.


1. Walker DI, McQuillan J, Taiwo M, Parks R, Stenton CA, Morgan H, et al. A highly specific Escherichia coli qPCR and its comparison with existing methods for environmental waters. Water Res. 2017;126: 101–110. doi:10.1016/j.watres.2017.08.032
2. Lenski RE, Rose MR, Simpson SC, Tadler SC. Long-Term Experimental Evolution in Escherichia coli . I . Adaptation and Divergence During 2000 Generations. 1991;138: 1315–1341. Available: www.jstor.org/stable/2462549
