## Supplementary figures and images for "Resident microbial communities inhibit growth and antibiotic resistance evolution of *Escherichia coli* in human gut microbiome samples"

### Supplemental Figure

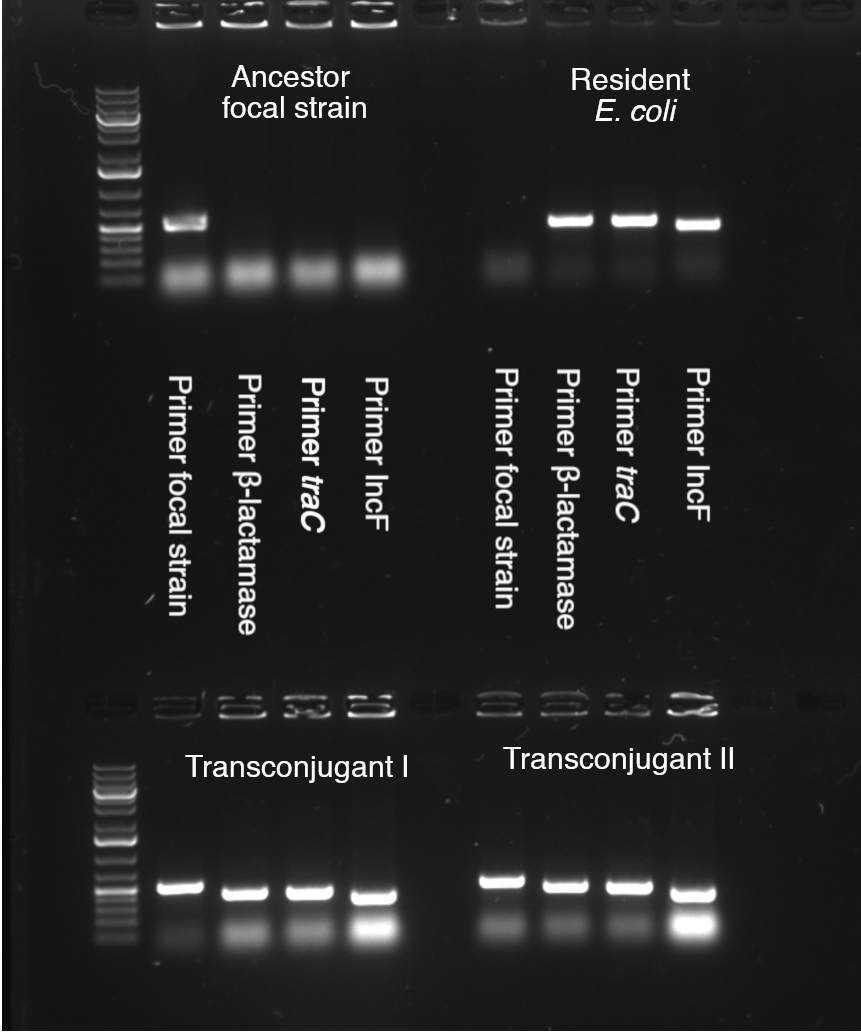

### Supplemental Figure 1

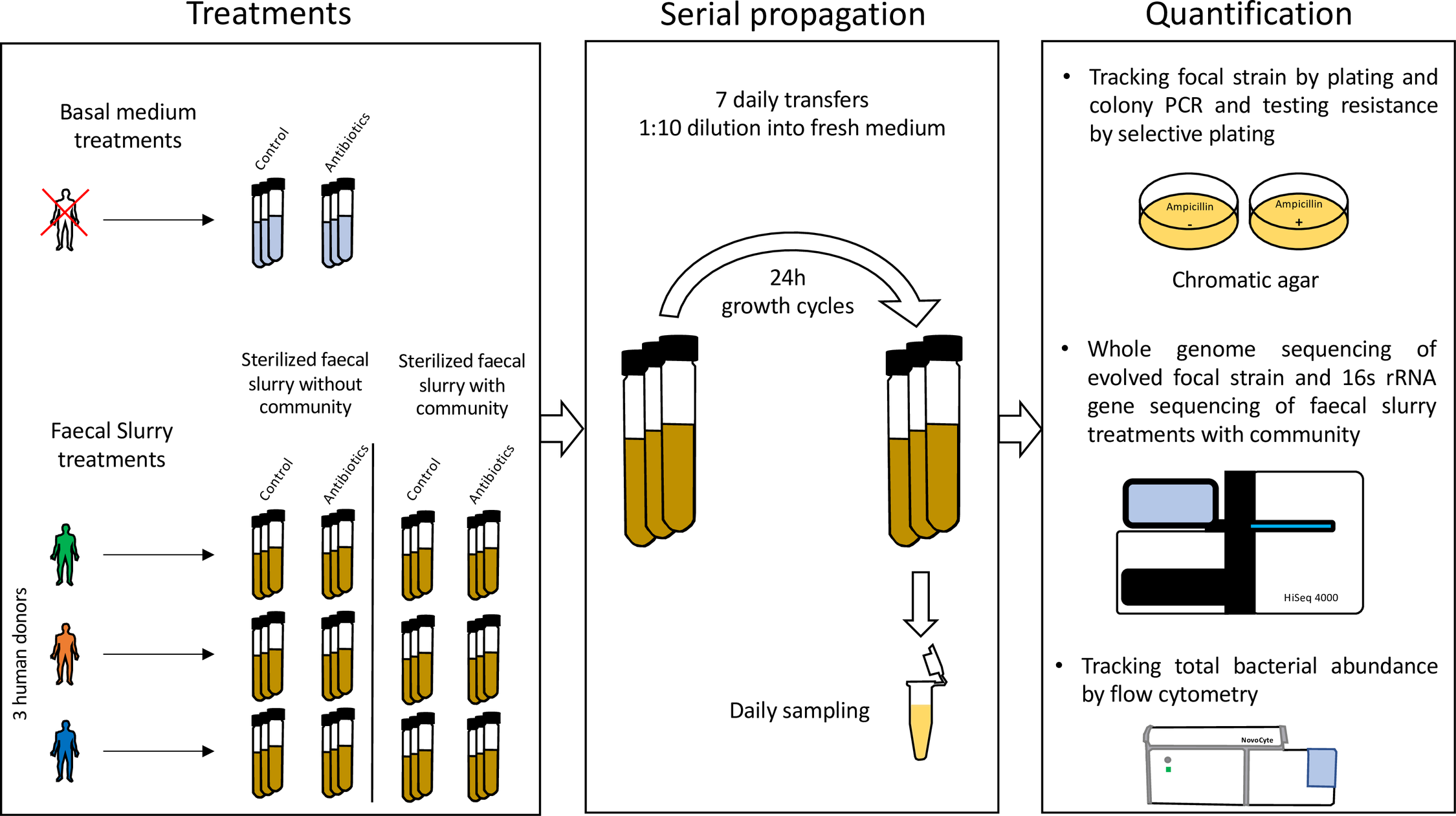

### Supplemental Figure 2

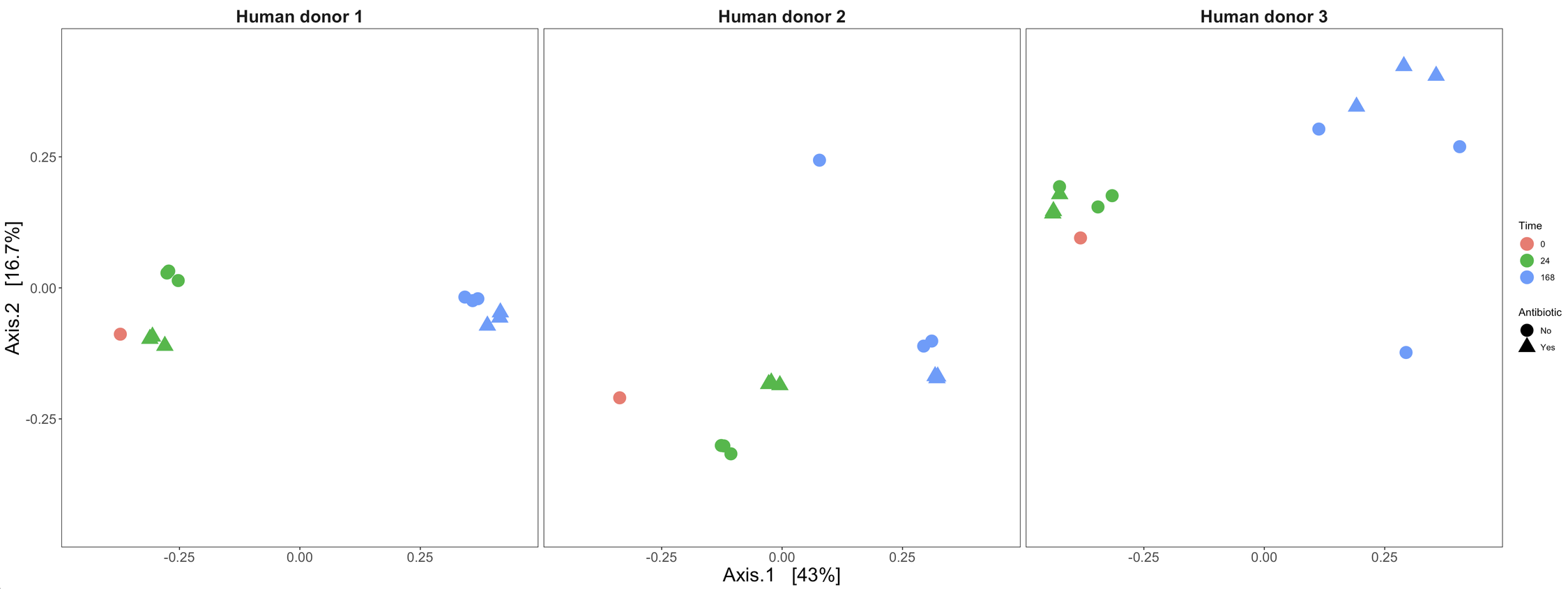

### Supplemental Figure 3

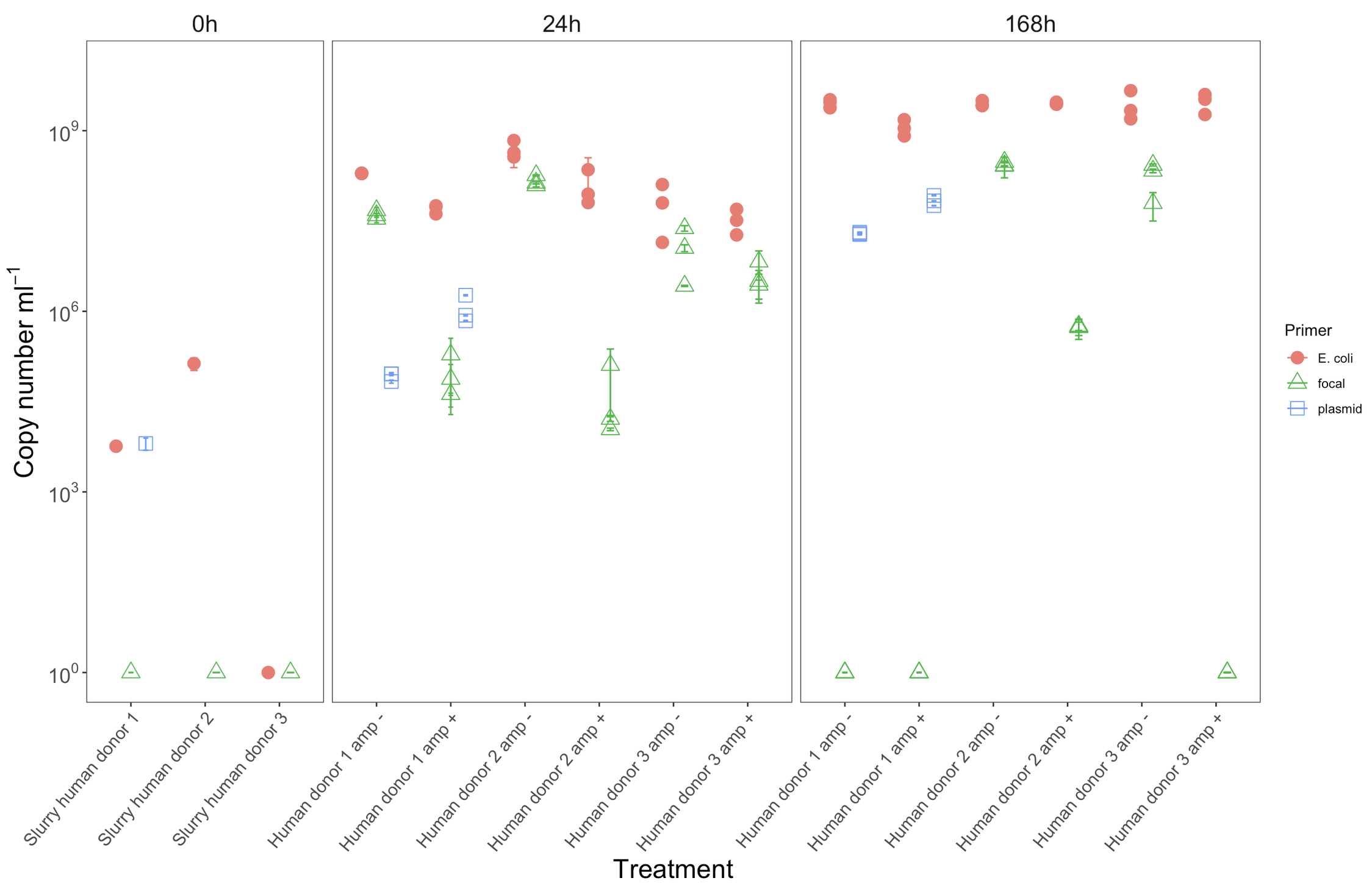

### Supplemental Figure 5

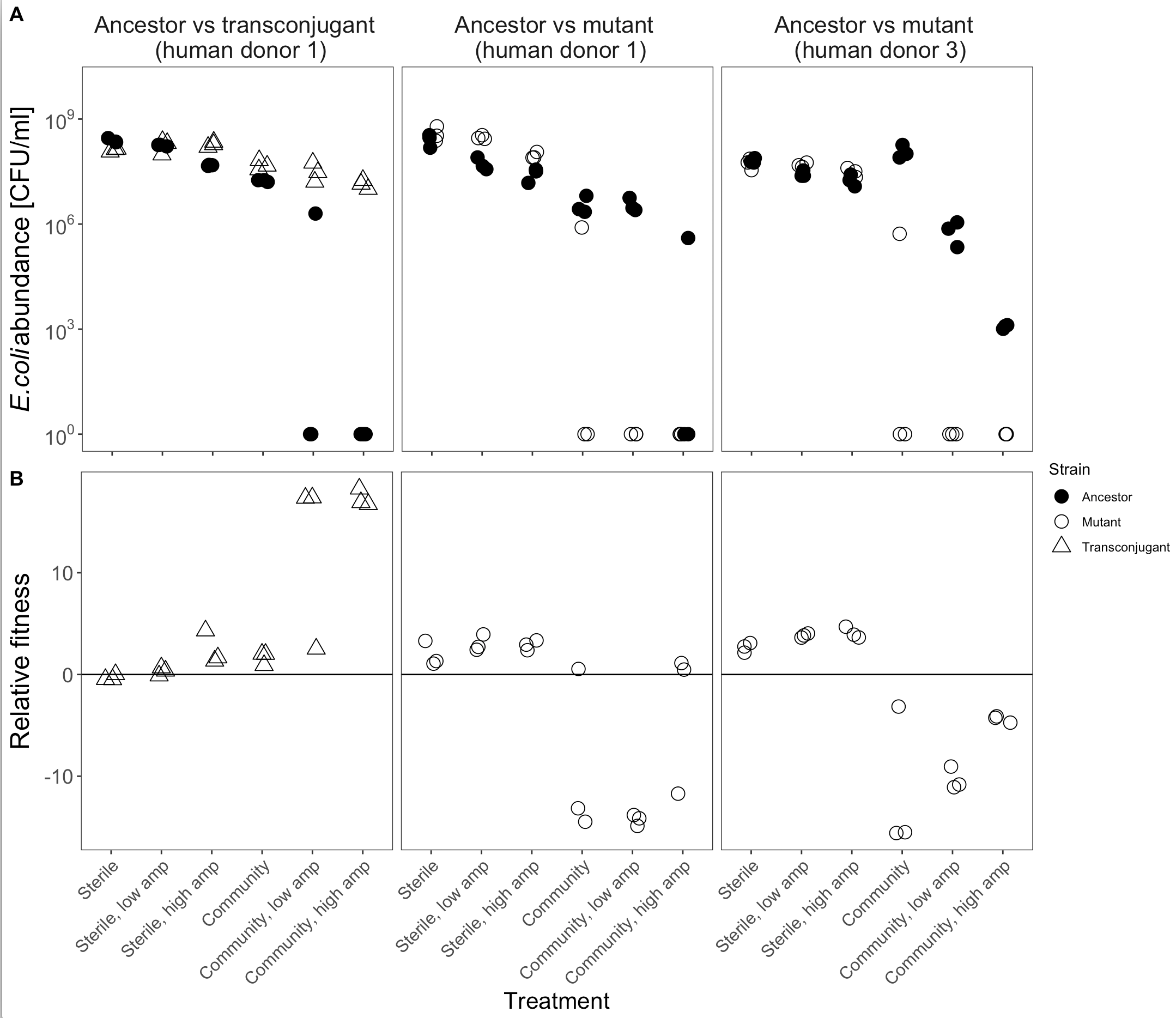

### Supplemental Figure 6

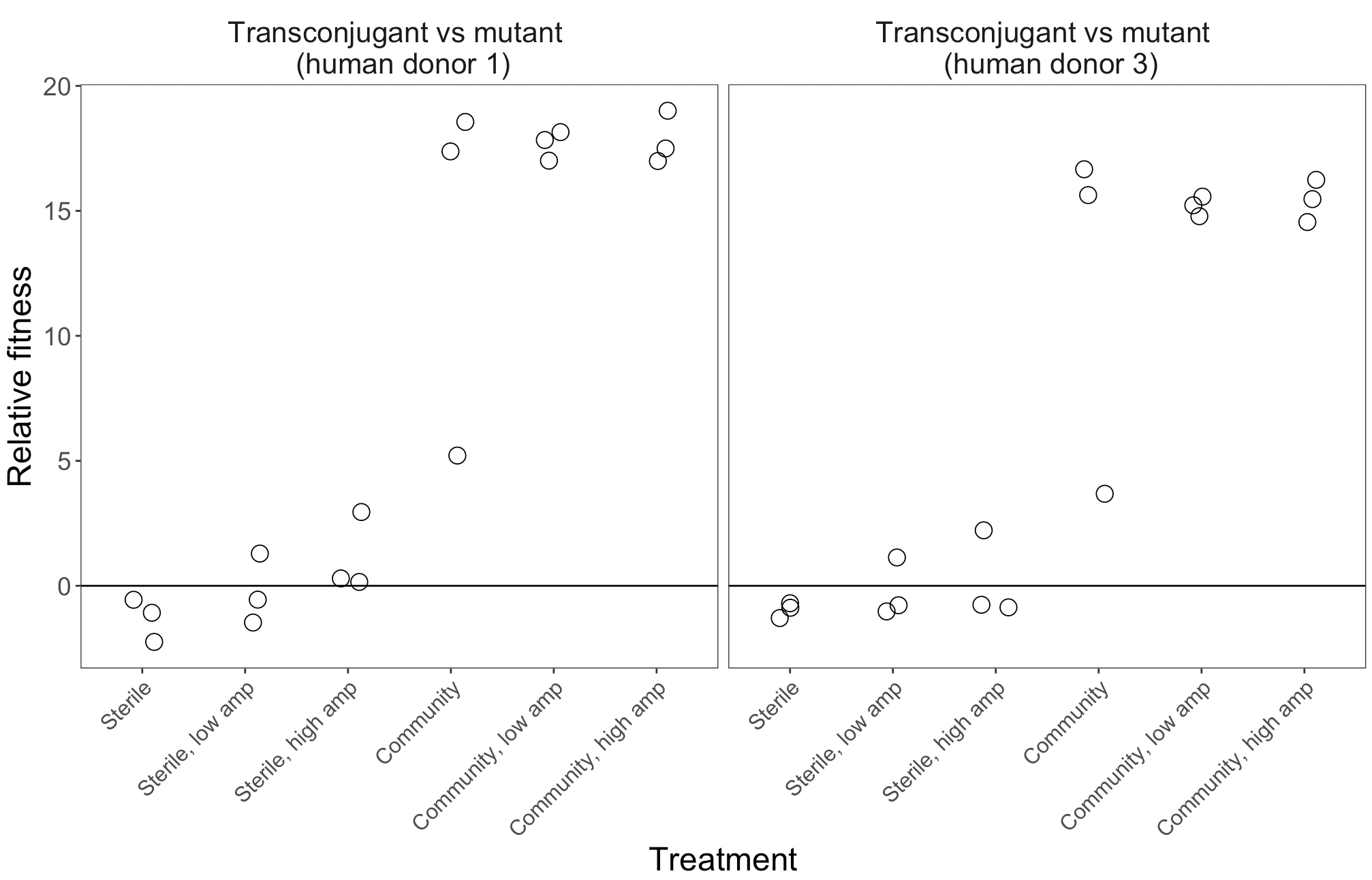

### Supplemental Figure 7

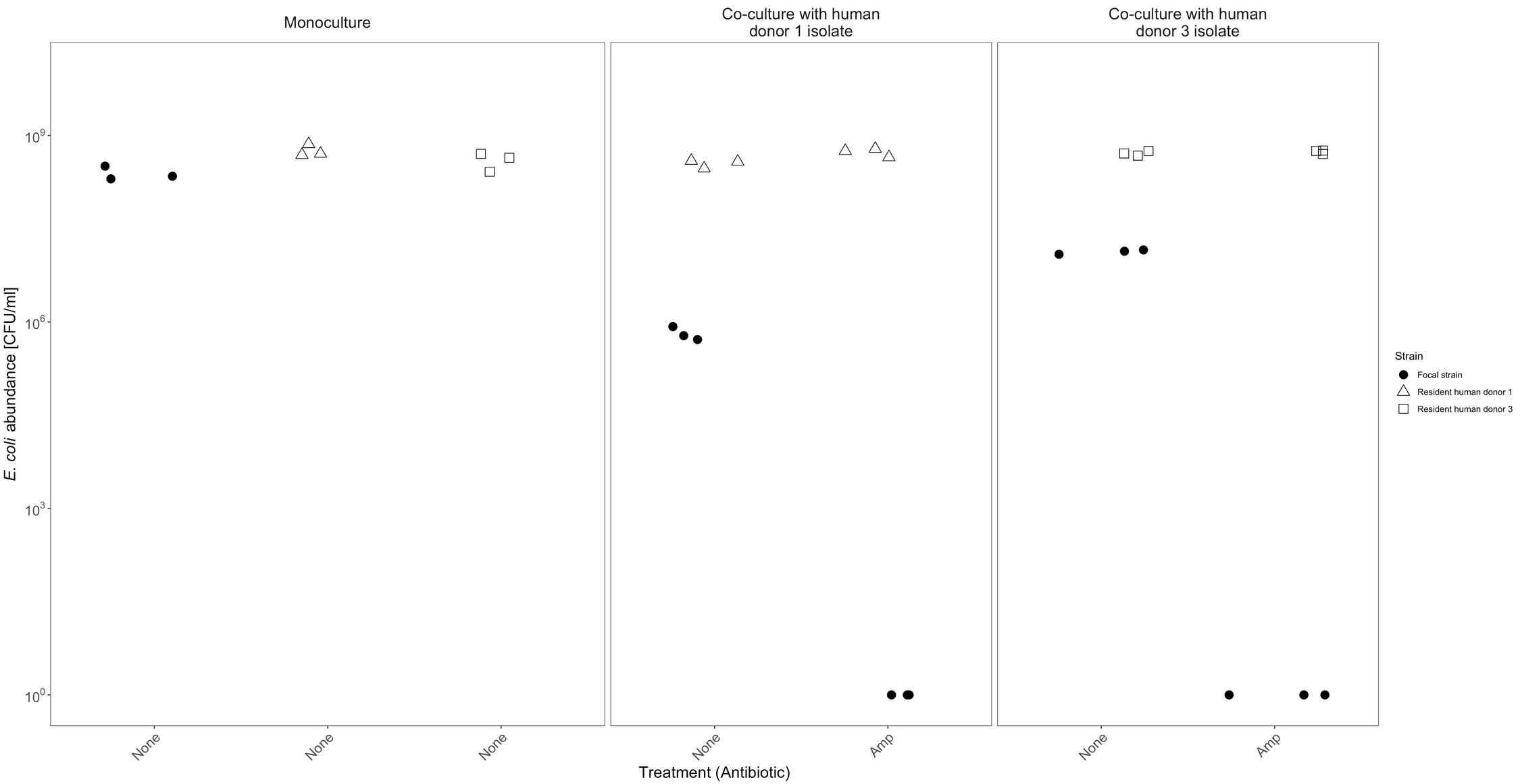

### Supplemental Figure 8

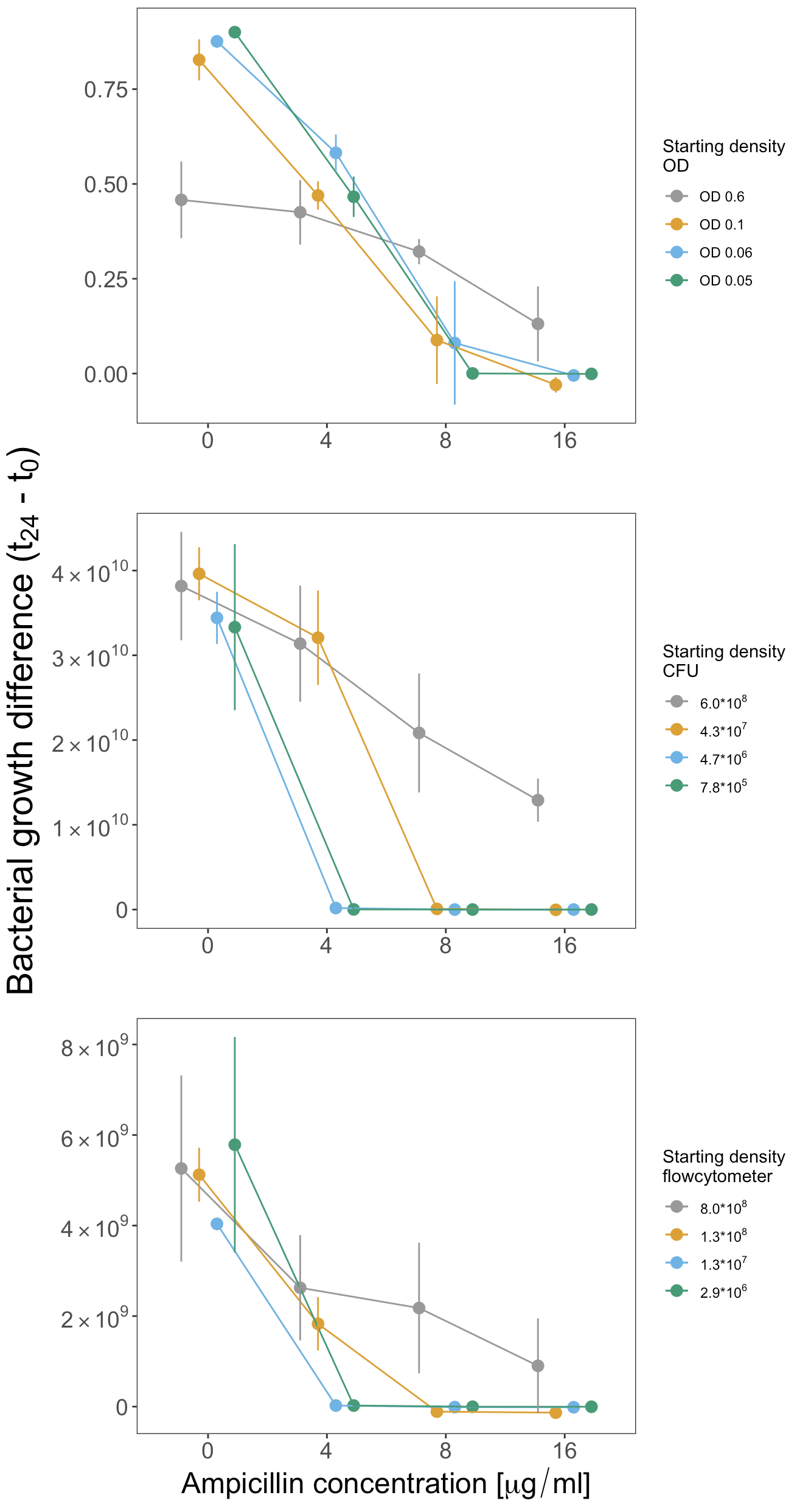
