## Supplementary material for "Resident microbial communities inhibit growth and antibiotic resistance evolution of *Escherichia coli* in human gut microbiome samples": S1 Model

### S1 model of plasmid transfer and transconjugant growth

The plasmid we identified in the resident microbial community of Human Donor 1 was conjugative on agar, but not in our gut microcosm system (Fig 5B). We were interested in whether a hypothetical plasmid with similar properties, but that *is* conjugatively transferable in our gut microcosm system, would have resulted in a high frequency of transconjugants of the focal strain in our experiment. We therefore used a modified version of the model described by Simonsen et al. [1] extended by Huisman et al. (*manuscript in preparation; preprint expected Jan. 2020*), to simulate transconjugant abundances over time.

We modelled abundances of plasmid Donors (*D*, resident *E. coli* carrying the plasmid), Recipients (*R*, the focal strain) and Transconjugants (*T*) over 24h. We specified a different maximum growth rate ($\psi_{max}$) for *D* and *R*. The population dynamics for each strain depend on $\psi_{max}$ and the maximum plasmid transfer rate ($\gamma_{max}$), both as a function of resource concentration (*C*, expressed relative to a half saturation constant *q*, and with a derivative set to zero when $C \leq 0$):

$$\frac{dD}{dt} = \psi_D(C)D$$

$$\frac{dR}{dt} = \psi_R(C)R - \gamma(C)(T + D)R$$

$$\frac{dT}{dt} = \psi_T(C)T + \gamma(C)(T + D)R$$

$$\frac{dC}{dt} = -(\psi_D(C)D + \psi_R(C)R + \psi_T(C)T)$$

where

$$\psi_{D/R/T}(C) = \psi_{max}^{D/R/T} \frac{C}{C + q}$$

and

$$\gamma(C) = \gamma_{max} \frac{C}{C + q}$$

To obtain biologically plausible estimates of the transfer rate $\gamma_{max}$ for our hypothetical plasmid, we searched *Web of Science* for publications citing Simonsen *et al.* [1] (this model has been widely used to estimate transfer rates experimentally). We then searched within these 81 publications for "*Escherichia coli*". Of the remaining 61, we included only publications reporting experimental transfer rate estimates for *E. coli* in the same units / using the same model as Simonsen *et al.* [1]. We then retrieved the minimum and maximum estimate from each of these 15 papers [1-15] (panel A in the figure below). In simulations, we used values of $\gamma_{max}$ ranging approximately from the first quartile of the minima ($10^{-14}$) to the third quartile of the maxima

($10^{-9}$). This is not intended as an exclusive or definitive report of *E. coli* transfer rates, but gives us a starting point for the simulations based on empirical estimates.

For the other parameters and starting values, we used estimates that reflected our experimental data for the Human Donor 1 treatment in the presence of the community and ampicillin (Fig 1, S3 and S5). Thus, initial abundances of *D* and *R* ($10^4$ and $10^6$) were similar to those estimated by qPCR and plating. We simulated two phases: 2h without ampicillin, then 22h with ampicillin (as in our main experiment). For the ampicillin-free phase we assigned a growth rate for each strain of 1.15 (calculated as ln($N_{final}$ / $N_{initial}$) / *time in hours*)). In the presence of ampicillin we assigned growth rates $\psi_{max}^{D} = 0.1$; $\psi_{max}^{T} = 0.1$; $\psi_{max}^{R} = -0.3$. The relatively small growth rate for the donor in the presence vs absence of ampicillin is because the second growth phase is longer than the first, so the net change in abundance per hour is smaller. We took *C* as $10^{12}$ and *q* as $10^2$. This resulted in similar population dynamics (panel B) as we inferred from plating and qPCR in the main experiment and in competitions between transconjugants and the focal strain.

We compared the abundance of transconjugants (*T*) after 24h for various conjugation rates (panel C). This revealed we would only expect more than ~100 transconjugants per ml at relatively high transfer rates (>$10^{-11}$). Thus, simulations indicate we would have sampled transconjugants of the focal strain that had acquired resistance from the resident microbiota within 24h, but only if the transfer rates were relatively high. We emphasize the model makes several simplifying assumptions. We therefore interpret these simulations as an indication of plausibility, rather than a firm prediction.

As a final step, we also simulated the same scenario, but where the focal strain can grow in the presence of ampicillin at a similar rate as observed in the absence of the community in our main experiment ($\psi_{max}^{D} = 0.02$ in the second phase of the simulation). That is, here we model the abundance of transconjugants as above, but we remove the suppressive effect of resident microbiota on growth of the focal strain in the presence of antibiotics which we observed in the main experiment (Fig 1). Transconjugant abundances here were ~8 times higher than those where the community suppresses growth of the focal strain (panel C). This suggests the growth-suppressive effects we observed in our main experiment would reduce the likelihood of conjugative plasmids spreading in the invading focal strain.

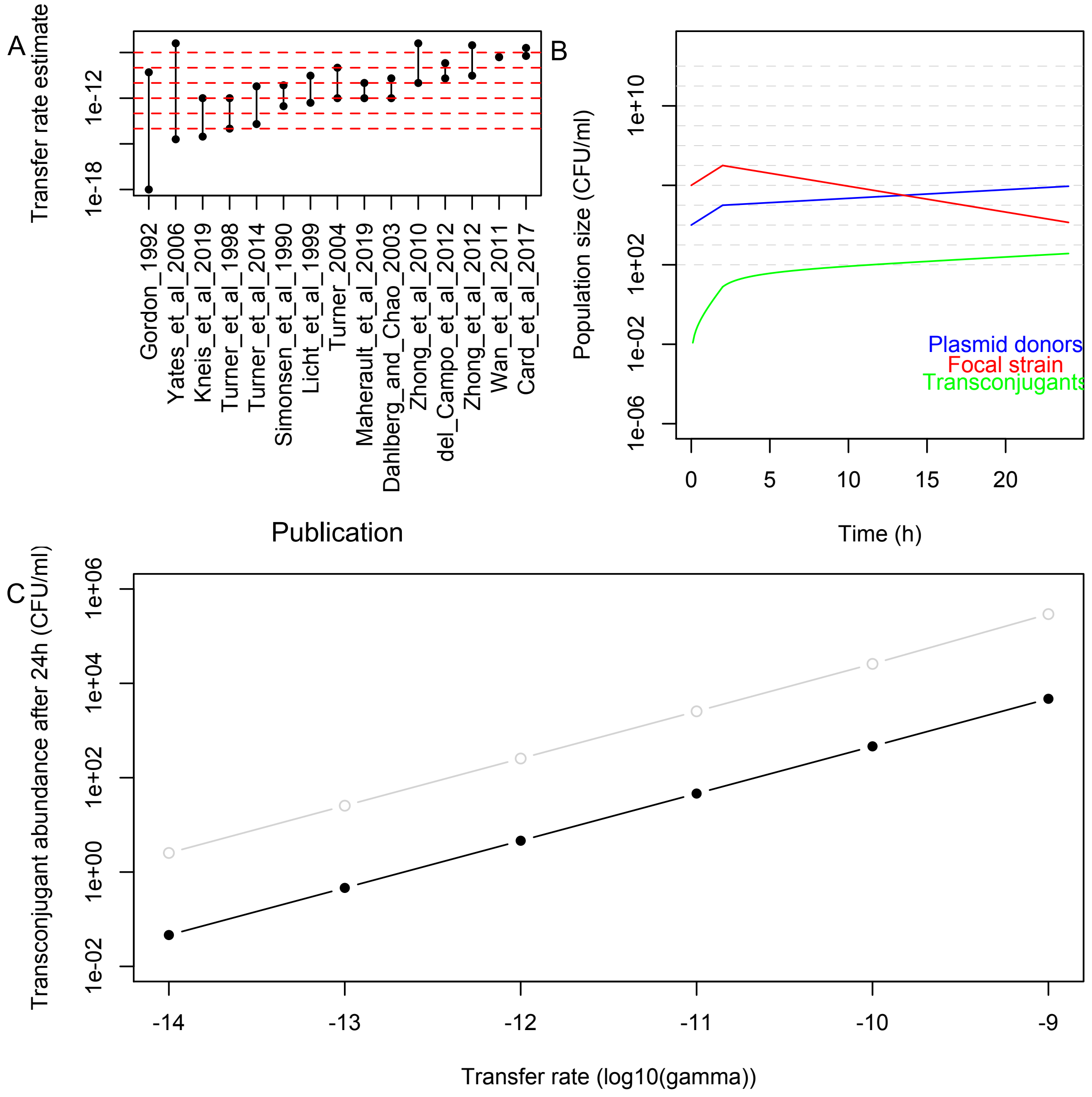


**Simulation of transfer dynamics for a hypothetical plasmid that is conjugative in our gut microcosm system and has similar fitness benefits to the plasmid we identified.** (A) Experimental transfer rate estimates from 15 publications using the same type of model we used in our simulations. (B) An example of the population dynamics in our simulations, with two phases (2h without ampicillin followed by 22h with ampicillin). These data are from a simulation with $\gamma_{max} = 10^{-11}$ (other parameters as described in the text above). (C) Abundance of transconjugants after 24h in simulations with various transfer rates. The black line is for the scenario described in the text above; the grey line shows the same scenario but where the resident microbiota no longer suppress growth of the focal strain (final paragraph of the above text).
