## Supplemental Table 1 for "Resident microbial communities inhibit growth and antibiotic resistance evolution of *Escherichia coli* in human gut microbiome samples"

**S1 Table: Abundance of the focal *E. coli* strain in treatments with and without ampicillin after 24h and averaged over the entire experiment.** *Treatment* indicates the different experimental groups (Basal is basal medium only; -Comm is sterilized faecal slurry; +Comm is "live" faecal slurry; -Amp and +Amp are with/without ampicillin). *Human donor* gives the human donor that faecal samples used in each treatment came from. $CFU_{24h}$ gives the mean and standard deviation of focal strain abundance after 24h. $Reduction_{24h}$ gives the average reduction in the presence of ampicillin relative to the absence of ampicillin for each combination of human donor and community. $CFU_{expt.}$ and $Reduction_{expt.}$ give the same but averaged across the entire seven-day experiment.

| **Treatment** | **Human donor** | **$CFU_{24h}$ (mean ±s.d.)** | | **$Reduction_{24h}$ (%)** | **$CFU_{expt.}$ (mean ±s.d.)** | | **$Reduction_{expt.}$ (%)** |
|---|---|---|---|---|---|---|---|
| Basal -Amp | None | $4.46\times10^7$ | $\pm6.07\times10^6$ | | $8.94\times10^7$ | $\pm3.85\times10^7$ | |
| Basal +Amp | None | $1.38\times10^7$ | $\pm2.68\times10^6$ | 69.028 | $3.54\times10^7$ | $\pm2.18\times10^7$ | 60.350 |
| -Comm -Amp | 1 | $1.33\times10^8$ | $\pm7.21\times10^6$ | | $1.35\times10^8$ | $\pm6.58\times10^7$ | |
| -Comm +Amp | 1 | $1.34\times10^7$ | $\pm2.91\times10^6$ | 89.955 | $3.03\times10^7$ | $\pm1.97\times10^7$ | 77.476 |
| -Comm -Amp | 2 | $1.58\times10^8$ | $\pm3.22\times10^7$ | | $1.43\times10^8$ | $\pm7.30\times10^7$ | |
| -Comm +Amp | 2 | $3.22\times10^7$ | $\pm6.52\times10^6$ | 79.594 | $4.75\times10^7$ | $\pm2.21\times10^7$ | 66.725 |
| -Comm -Amp | 3 | $2.04\times10^8$ | $\pm2.95\times10^7$ | | $1.24\times10^8$ | $\pm6.78\times10^7$ | |
| -Comm +Amp | 3 | $4.43\times10^7$ | $\pm8.59\times10^6$ | 78.322 | $3.27\times10^7$ | $\pm1.94\times10^7$ | 73.743 |
| +Comm -Amp | 1 | $6.41\times10^7$ | $\pm1.08\times10^7$ | | $2.30\times10^7$ | $\pm3.02\times10^7$ | |
| +Comm +Amp | 1 | $8.22\times10^3$ | $\pm1.02\times10^4$ | 99.987 | $1.17\times10^3$ | $\pm3.11\times10^3$ | 99.995 |
| +Comm -Amp | 2 | $9.25\times10^7$ | $\pm4.85\times10^7$ | | $1.09\times10^8$ | $\pm5.86\times10^7$ | |
| +Comm +Amp | 2 | $4.23\times10^3$ | $\pm4.84\times10^3$ | 99.995 | $7.30\times10^4$ | $\pm1.57\times10^5$ | 99.933 |
| +Comm -Amp | 3 | $4.40\times10^6$ | $\pm3.03\times10^6$ | | $5.74\times10^7$ | $\pm5.15\times10^7$ | |
| +Comm +Amp | 3 | $7.08\times10^5$ | $\pm3.60\times10^5$ | 83.909 | $1.31\times10^7$ | $\pm3.00\times10^7$ | 77.226 |
