## Supplemental Table 2 for "Resident microbial communities inhibit growth and antibiotic resistance evolution of *Escherichia coli* in human gut microbiome samples"

**S2 Table: Fraction of focal strain on total E.coli abundance determined by qPCR and a mixed calculation based of colony PCR, flow cytometry and amplicon data.** *Treatment* and *Human donor* indicate the same experimental groups as in Sup. table 1. *Time point* gives the time after the start of the experiment when the samples were isolated. Frequency of the focal strain compared to total *E. coli* abundance was either determined by qPCR with primers specific for focal strain and *E.coli* or a mixed calculation of abundance of focal strain based on colony PCR, total bacterial abundance measured by flow cytometry and total *E. coli frequency* based on amplicon data. *E.coli* frequency based on total *Enterobacteriaceae* abundance was determined by reads distribution of the amplicon sequencing results. Frequencies shown are mean values of three replicates and the standard deviation

| | | | Frequency of focal strain compared to total *E. coli* abundance based on | | *E.coli* frequency of total *Enterobacteriaceae* based on |
|---|---|---|---|---|---|
| **Treatment** | **Human donor** | **Time point [h]** | **qPCR (mean % ± s.d. )** | **Colony PCR/ Flow cytometer/ amplicon (mean % ± s.d. )** | **Amplicon (mean % ± s.d. )** |
| +Comm -Amp | 1 | 24 | 20.9 ± 3.1 | 66.9 ± 5.9 | 99.997 ± 0.003 |
| +Comm +Amp | 1 | 24 | 0.2 ± 0.1 | 0.1 ± 0.03 | 99.93 ± 0.007 |
| +Comm -Amp | 2 | 24 | 35.1 ± 11.1 | 27.9 ± 8.3 | 99.97 ± 0.02 |
| +Comm +Amp | 2 | 24 | 0.07 ± 0.1 | 0.004 ± 0.004 | 99.99 ± 0.006 |
| +Comm -Amp | 3 | 24 | 18.6 ± 0.6 | 16.6 ± 3.8 | 99.999 ± 0.001 |
| +Comm +Amp | 3 | 24 | 12.9 ± 2.6 | 8.6 ± 2.5 | 99.998 ± 0.004 |
| +Comm -Amp | 1 | 168 | ND | 0 | 100 ± 0.001 |
| +Comm +Amp | 1 | 168 | ND | 0 | 99.999 ± 0.001 |
| +Comm -Amp | 2 | 168 | 9.6 ± 0.1 | 32.9 ± 2.5 | 99.773 ± 0.2 |
| +Comm +Amp | 2 | 168 | 0.02 ± 0.001 | 0.0001± 0.0001 | 99.474 ± 0.64 |
| +Comm -Amp | 3 | 168 | 9.4± 7.0 | 23.3± 5.6 | 99.996 ± 0.007 |
| +Comm +Amp | 3 | 168 | ND | 0 | 99.992 ± 0.01 |
