## Supplemental Table 3 for "Resident microbial communities inhibit growth and antibiotic resistance evolution of *Escherichia coli* in human gut microbiome samples"

**S3 Table (continued on next page): Genomic variants found in randomly selected colony isolates of the focal strain picked from ampicillin-free agar plates at the end of the experiment.** *Treatment group* indicates if strain evolved in presence or absence of community (Com) or ampicillin (Amp). *Human donor* gives the origin of the faecal samples. Arrows indicate that intergenic region, list of genes indicate that several genes were affected by the same mutation and delta sign indicates larger deletion on the chromosome.

| Treatment group | Human donor | Replicate | *arpA <> iclR* | *casA <> ygcB* | *clsA* | *entD* | *frsA <> phoE* | *gatY <> fbaB* | *gatZ* | *gfvA+B+C, insA+B* | *gtrS* | *hsdS* | *insI1, insX* | *insN* | *intD* | *intD, xisD,exoD, peaD* | *iraM <> ycgX* | *mmuM+P, afuB+C,* | *ompF <> asnS* | *ompR* | *ompT* | *opgB* | *rpoD* | *serS* | *tfaD* | *tfaD <> ybcY* | *tfaX <> appY* | *waaS* | *waaU* | *wbbK* | *wecB* | *yaiO* | *yaiP* | *yaiS <> tauA* | *ybbD* | *ybcy* | *ybfK <> kdpE* | *yddK* | *ydjO* | *yedR <> yedS* | *yfcV <> sixA* | *yfjI* | *ygeF* | *yhhI* | *yibV* | *yjbS* | *yjgL* | *yjhB* | *yjiS <> yjiT* | Δ 566060-584707 | Δ 568024-586058 |
|---|---|---|---|---|---|---|---|---|---|---|---|---|---|---|---|---|---|---|---|---|---|---|---|---|---|---|---|---|---|---|---|---|---|---|---|---|---|---|---|---|---|---|---|---|---|---|---|---|---|---|
| Basal -Amp | None | 1 | | | | | | | | | | | | x | | | | | | | | | | | | | | | | | | | | | | | | | | | | | | | | | | | |
| | | 2 | | | | | | | | | | | ▲ | | | | | ▲ | | | | | | | | | | | | | | x | | | | | | | | | | | | | | | | | | | |
| | | 3 | No mutations detected | | | | | | | | | | | | | | | | | | | | | | | | | | | | | | | | | | | | | | | | | | | | | | | | |
| -Com -Amp | 1 | 1 | No mutations detected | | | | | | | | | | | | | | | | | | | | | | | | | | | | | | | | | | | | | | | | | | | | | | | | |
| | | 2 | | | | | | | | | | | | | | | | | | | | | | ● | | | | | | | | | | | | | | | | | | | | | | | | | | | |
| | | 3 | No mutations detected | | | | | | | | | | | | | | | | | | | | | | | | | | | | | | | | | | | | | | | | | | | | | | | | |
| Com -Amp | 1 | 1 | Focal strain below detection limit | | | | | | | | | | | | | | | | | | | | | | | | | | | | | | | | | | | | | | | | | | | | | | | | |
| | | 2 | | | | | | | | | | | | | | | | | | | | | | | | | | | | | | | | | | | | | | | | | | | | | | | | | |
| | | 3 | | | | | | | | | | | | | | | | | | | | | | | | | | | | | | | | | | | | | | | | | | | | | | | | | |
| -Com -Amp | 2 | 1 | No mutations detected | | | | | | | | | | | | | | | | | | | | | | | | | | | | | | | | | | | | | | | | | | | | | | | | |
| | | 2 | | | | | | | | | | | | | | | | | | | | | | | | | | | | | | | | | | | | | | | | | | | | | | | | | |
| | | 3 | | | | | | | | | ▲ | | | | | | | | | | | ● | | | | | | | | | | ● | | | | | | | | | | | | | | | | | | | |
| +Com -Amp | 2 | 1 | No mutations detected | | | | | | | | | | | | | | | | | | | | | | | | | | | | | | | | | | | | | | | | | | | | | | | | |
| | | 2 | | | | | | | | | | | | | | | | | | | | | | | | | | | | | | | | | | | | | | | | | | | | | | | | | |
| | | 3 | | | | | | | | | | | | | | | | | | | | | | | | | | | | | | | | | | | | | | | | | | | | | | | | | |
| -Com -Amp | 3 | 1 | | | | | | | | | | | | | | | | | | | | | | | | | | | | | | | | | | | | | | | | x | | | | | | | | | |
| | | 2 | | | | | | | | | ▲ | | | | | | | | | | | | | | | | | | ▲ | | | | | ▲ | | | | | | | | | | | x | | | | | | |
| | | 3 | No mutations detected | | | | | | | | | | | | | | | | | | | | | | | | | | | | | | | | | | | | | | | | | | | | | | | | |
| +Com -Amp | 3 | 1 | | | | | | | | x | | | | | ● | | | | | | ● | | | | ● | ● | | | | | | | | | | ● | | | | | | | | | | | | | | | ▲ |
| | | 2 | | | | | | | | | | | | x | | | | | | | | | | | | | | | | | | | | | | | | | | | | | | | | | | | | | |
| | | 3 | | | | x | ● | | | | | | | | | ▲ | | | | | | | | | | | ▲ | | | | | | x | | | | | ● | | | | | | | | | | | | | ▲ |

▲ Deletion x Insertion ● SNP

**Continuation S3 Table: Genomic variants found in randomly selected colony isolates of the focal strain picked from ampicillin-free agar plates at the end of the experiment.**

| Treatment group | Human donor | Replicate | arpA <> iclR | casA <> ygcB | clsA | entD | frsA <> phoE | gatY <> fbaB | gatZ | gfvA+B+C, insA+B | gtrS | hsdS | insI1, insX | insN | intD | intD, xisD,exoD, | iraM <> ycgX | mmuM+P, afuB+C, | ompF <> asnS | ompR | ompT | opgB | rpoD | serS | tfaD | tfaD <> ybcY | tfaX <> appY | waaS | waaU | wbbK | wecB | yaiO | yaiP | yaiS <> tauA | ybbD | ybcy | ybfK <> kdpE | yddK | ydjO | yedR <> yedS | yfcV <> sixA | yfjl | ygeF | yhhI | yibV | yjbS | yjgL | yjhB | yjiS <> yjiT | Δ 566060-584707 | Δ 568024-586058 |
|---|---|---|---|---|---|---|---|---|---|---|---|---|---|---|---|---|---|---|---|---|---|---|---|---|---|---|---|---|---|---|---|---|---|---|---|---|---|---|---|---|---|---|---|---|---|---|---|---|---|---|---|
| Basal +Amp | None | 1 | | | | | | | | | | | | x | | | | | | | | | | | | | | | | | | | | | | | | | | | | | | | | | | | |
| | | 2 | No mutations detected | | | | | | | | | | | | | | | | | | | | | | | | | | | | | | | | | | | | | | | | | | | | | | | |
| | | 3 | | | | | | | | | | | | | | | | | | | | | | | | | | | | | | | | | | | | | | | | | | | | | | | |
| -Com +Amp | 1 | 1 | | | | | | | | | | | | | | | | | | | | | ● | | | | | | | | | | | | ▲ | | | | | ▲ | | x | | | | | | | | | |
| | | 2 | No mutations detected | | | | | | | | | | | | | | | | | | | | | | | | | | | | | | | | | | | | | | | | | | | | | | | |
| | | 3 | | | | | | | | | | | | x | | | | | | ● | | | | | | | | | | | | | | | | | | | | | | | | | | | | | | | |
| +Com +Amp | 1 | 1 | | | | | | | | | | | | | | | | | | | | | | | | | | | | | | | | | | | | | | | | | | | | | | | |
| | | 2 | Focal strain below detection limit | | | | | | | | | | | | | | | | | | | | | | | | | | | | | | | | | | | | | | | | | | | | | | | |
| | | 3 | | | | | | | | | | | | | | | | | | | | | | | | | | | | | | | | | | | | | | | | | | | | | | | |
| -Com +Amp | 2 | 1 | | | | | | | | | ▲ | | | | | | | | x | | | | | | | | | | | | | | | | | | | | | | | | | | | | | | | | |
| | | 2 | ▲ | | x | | | | | | | | | | | | | | | ● | | | | | | | | | | | | | | | | | | | | | | | | ▲ | | | ▲ | | | |
| | | 3 | | | | | | ● | | | | | | | | | | | | ● | | | | | | | ▲ | | | | | | | | | | | | | | | | | | | | | | | | |
| +Com +Amp | 2 | 1 | | | | | | | | | | | | | | | | | | | | | | | | | | | | | | | | | | | | | | | | | | | | | | | |
| | | 2 | | ▲ | | | | | | | | | | | | | ▲ | | | | | | | | | | | ▲ | ▲ | ▲ | | | | | | | ▲ | | ▲ | | ▲ | | ▲ | | | ▲ | ▲ | | | | |
| | | 3 | | | | | | | | | ▲ | | | | | | | | | | | | | | | | | | | | | | | | | | | | | | | | | | | | | | ▲ | | |
| -Com +Amp | 3 | 1 | | | | | | | | | | | | x | | | | | | | | | | | | | | | | | | | | | | | | | | | | | | | | | ▲ | | | |
| | | 2 | No mutations detected | | | | | | | | | | | | | | | | | | | | | | | | | | | | | | | | | | | | | | | | | | | | | | | |
| | | 3 | | | | | | | | | | | | | | | | | | | | | | | | | | | | | | | | | | | | | | | | | | | | | | | |
| +Com +Amp | 3 | 1 | | | | | | | | | | | | | | | | | | | | | | | | | | | | | | | | | | | | | | | | | | | | | | | |
| | | 2 | Focal strain below detection limit | | | | | | | | | | | | | | | | | | | | | | | | | | | | | | | | | | | | | | | | | | | | | | | |
| | | 3 | | | | | | | | | | | | | | | | | | | | | | | | | | | | | | | | | | | | | | | | | | | | | | | |

▲ Deletion x Insertion ● SNP
