## Supplemental Table 4 for "Resident microbial communities inhibit growth and antibiotic resistance evolution of *Escherichia coli* in human gut microbiome samples"

**S4 Table: Antibiotic resistance genes, plasmid replicons, genes involved in conjugative transfer and formation of type 6 secretion system found on plasmid 1 of isolate from human donor 1 resident *E. coli* community and on the chromosome of human donor 3 resident *E.*coli isolates of each replicate population**

| | | Plasmid 1 from human donor 1 without antibiotics of replicate community: | | | Plasmid 1 from human donor 1 with antibiotics of replicate community: | | | Chromosome from human donor 3 without antibiotics of replicate community: | | | Chromosome from human donor 3 with antibiotics of replicate community: | | |
|---|---|---|---|---|---|---|---|---|---|---|---|---|---|
| **Genomic feature** | **Function** | 1 | 2 | 3 | 1 | 2 | 3 | 1 | 2 | 3 | 1 | 2 | 3 |
| blaTem-1b | Beta lactam resistance | ✓ | ✓ | ✓ | ✓ | ✓ | ✓ | ✓ | ✓ | ✓ | ✓ | ✓ | ✓ |
| aph(3) | Aminoglycoside resistance | ✓ | ✓ | ✓ | ✓ | ✓ | ✓ | ✓ | ✓ | ✓ | ✓ | ✓ | ✓ |
| aph(6) | Aminoglycoside resistance | ✓ | ✓ | ✓ | ✓ | ✓ | ✓ | ✓ | ✓ | ✓ | ✓ | ✓ | ✓ |
| sul2 | Sulphonamide resistance | ✓ | ✓ | ✓ | ✓ | ✓ | ✓ | ✓ | ✓ | ✓ | ✓ | ✓ | ✓ |
| tet(A) | Tetracycline resistance | ✕ | ✕ | ✕ | ✕ | ✕ | ✕ | ✓ | ✓ | ✓ | ✓ | ✓ | ✓ |
| IncQ1 | Replicon | ✕ | ✕ | ✕ | ✕ | ✕ | ✕ | ✓ | ✓ | ✓ | ✓ | ✓ | ✓ |
| IncFIC(FII) | Replicon | ✓ | ✓ | ✓ | ✓ | ✓ | ✓ | ✕ | ✕ | ✕ | ✕ | ✕ | ✕ |
| IncFIA | Replicon | ✓ | ✓ | ✓ | ✓ | ✓ | ✓ | ✕ | ✕ | ✕ | ✕ | ✕ | ✕ |
| IncFIB | Replicon | ✓ | ✓ | ✓ | ✓ | ✓ | ✓ | ✕ | ✕ | ✕ | ✕ | ✕ | ✕ |
| tra genes [a] | Involved in conjugative transfer | ✓ | ✓ | ✓ | ✓ | ✓ | ✓ | ✕ | ✕ | ✕ | ✕ | ✕ | ✕ |
| tss genes [a] | Type 6 secretion system | ✓ | ✓ | ✓ | ✓ | ✓ | ✓ | ✓ | ✓ | ✓ | ✓ | ✓ | ✓ |

[a] For names of involved genes see Fig. 4
