## Supplemental Table 5 for "Resident microbial communities inhibit growth and antibiotic resistance evolution of *Escherichia coli* in human gut microbiome samples"

**S5 Table: IC90 values of ancestor and ampicillin resistant evolved strains.** IC90 was defined as the concentration where the OD value was less than 10% of the untreated control.

| Isolate | Human donor | Isolated from Replicate population | IC90 concentration [µg/ml] |
|---|---|---|---|
| Ancestor focal strain | None | None | 8 |
| Basal +Amp | None | 1 | 20 |
| Basal +Amp | None | 2 | 26 |
| Basal +Amp | None | 3 | 16 |
| -Com +Amp | 1 | 1 | 20 |
| -Com +Amp | 1 | 2 | 16 |
| -Com +Amp | 1 | 3 | 16 |
| -Com +Amp | 3 | 1 | 20 |
| -Com +Amp | 3 | 2 | 20 |
| Ancestor transconjugant | None | None | 8 |
| Transconjugant | None | None | >60 |
| Resident E. coli | 1 | None | >60 |
