## Supplemental Table 6 for "Resident microbial communities inhibit growth and antibiotic resistance evolution of *Escherichia coli* in human gut microbiome samples"

**S6 Table A: Assembly statistics for genome sequencing on Illumina platform of resident *E. coli* isolated from the resident microbiota of human donors 1 and 3.** *Ampicillin treatment*, *Human donor* and *Replicate microcosm* indicate the treatment group of the main experiment each isolate was taken from before sequencing.

| Ampicillin treatment | Human donor | Replicate microcosm | No. contigs | Total genome length [bp] | Largest contig [bp] | N50 | L50 | GC (%) |
|---|---|---|---|---|---|---|---|---|
| -Amp | 1 | 1 | 88 | 5214174 | 557565 | 207210 | 8 | 50.5 |
| -Amp | 1 | 2 | 103 | 5185519 | 334964 | 153591 | 12 | 50.5 |
| -Amp | 1 | 3 | 115 | 5207591 | 346088 | 158712 | 12 | 50.5 |
| +Amp | 1 | 1 | 4 | 5330642 | 5158979 | 5158979 | 1 | 50.5 |
| +Amp | 1 | 1 | 86 | 5218659 | 522040 | 196713 | 8 | 50.5 |
| +Amp | 1 | 2 | 99 | 5207712 | 522040 | 162556 | 10 | 50.5 |
| +Amp | 1 | 3 | 91 | 5216794 | 520971 | 189113 | 9 | 50.5 |
| -Amp | 3 | 1 | 95 | 5095638 | 570677 | 191144 | 9 | 50.6 |
| -Amp | 3 | 2 | 110 | 5094217 | 570677 | 191168 | 9 | 50.6 |
| -Amp | 3 | 3 | 86 | 5096162 | 540821 | 218238 | 8 | 50.6 |
| +Amp | 3 | 1 | 4 | 5186270 | 5072055 | 5072055 | 1 | 50.6 |
| +Amp | 3 | 1 | 105 | 5100479 | 373745 | 203252 | 10 | 50.6 |
| +Amp | 3 | 2 | 93 | 5093766 | 570638 | 191168 | 9 | 50.6 |
| +Amp | 3 | 3 | 97 | 5099961 | 570542 | 203252 | 9 | 50.6 |

**S6 Table B: Assembly statistics for genome sequencing on MinION platform of resident *E. coli* isolated from the resident microbiota of human donors 1 and 3.** *Ampicillin treatment*, *Human donor* and *Replicate microcosm* indicate the treatment group of the main experiment each isolate was taken from before sequencing and *Circular* indicates if the contig was closed or not.

| **Ampicillin treatment** | **Human donor** | **Replicate microcosm** | **Contig 1** | | **Contig 2** | | **Contig 3** | | **Contig 4** | | **Contig 5** | |
|---|---|---|---|---|---|---|---|---|---|---|---|---|
| | | | Length [bp] | Circular | Length [bp] | Circular | Length [bp] | Circular | Length [bp] | Circular | Length [bp] | Circular |
| +Amp | 1 | 1 | 5158979 | Yes | 163562 | Yes | 6647 | Yes | 1454 | No | - | - |
| +Amp | 3 | 1 | 5072055 | No | 109229 | No | 3030 | No | 1584 | No | 372 | No |
