## Supplemental Table 7 for "Resident microbial communities inhibit growth and antibiotic resistance evolution of *Escherichia coli* in human gut microbiome samples"

**S7 Table: List of all sequenced isolates:** *Treatment, Human donor* and *Replicate population* indicates where the strain was isolated from. Strains were isolated either from ampicillin-free or ampicillin containing plates. *Strain* indicates if it is the focal or resident strain. *Sample ID* gives the short name used for sequencing and *Platform* indicates if the isolate was sequenced with Illumnia or Minion. *ENA accession* gives the accession number of the isolate in the repository and *Ampicillin resistant* indicates if the isolate could grow at the MIC of the ancestor strain.

| Treatment | Human donor | Replicate population | Isolated from | Strain | Sample ID | Platform | ENA Accession | Ampicillin resistant |
|---|---|---|---|---|---|---|---|---|
| Basal -Amp | none | 1 | Ampicillin-free plates | Focal | Strep_B1 | Illumina | ERS4256211 | No |
| Basal -Amp | none | 2 | Ampicillin-free plates | Focal | Strep_B2 | Illumina | ERS4256212 | No |
| Basal -Amp | none | 3 | Ampicillin-free plates | Focal | Strep_B3 | Illumina | ERS4256213 | No |
| Basal +Amp | none | 1 | Ampicillin-free plates | Focal | Strep_BA1 | Illumina | ERS4256214 | No |
| Basal +Amp | none | 2 | Ampicillin-free plates | Focal | Strep_BA2 | Illumina | ERS4256215 | No |
| Basal +Amp | none | 3 | Ampicillin-free plates | Focal | Strep_BA3 | Illumina | ERS4256216 | No |
| -Com -Amp | 1 | 1 | Ampicillin-free plates | Focal | Strep_D1S1 | Illumina | ERS4256217 | No |
| -Com -Amp | 1 | 2 | Ampicillin-free plates | Focal | Strep_D1S2 | Illumina | ERS4256218 | No |
| -Com -Amp | 1 | 3 | Ampicillin-free plates | Focal | Strep_D1S3 | Illumina | ERS4256219 | No |
| -Com +Amp | 1 | 1 | Ampicillin-free plates | Focal | Strep_D1SA1 | Illumina | ERS4256220 | Yes |
| -Com +Amp | 1 | 2 | Ampicillin-free plates | Focal | Strep_D1SA2 | Illumina | ERS4256221 | No |
| -Com +Amp | 1 | 3 | Ampicillin-free plates | Focal | Strep_D1SA3 | Illumina | ERS4256222 | No |
| -Com -Amp | 2 | 1 | Ampicillin-free plates | Focal | Strep_D2S1 | Illumina | ERS4256229 | No |
| -Com -Amp | 2 | 2 | Ampicillin-free plates | Focal | Strep_D2S2 | Illumina | ERS4256230 | No |
| -Com -Amp | 2 | 3 | Ampicillin-free plates | Focal | Strep_D2S3 | Illumina | ERS4256231 | No |
| -Com +Amp | 2 | 1 | Ampicillin-free plates | Focal | Strep_D2SA1 | Illumina | ERS4256232 | No |
| -Com +Amp | 2 | 2 | Ampicillin-free plates | Focal | Strep_D2SA2 | Illumina | ERS4256233 | No |
| -Com +Amp | 2 | 3 | Ampicillin-free plates | Focal | Strep_D2SA3 | Illumina | ERS4256234 | No |
| +Com -Amp | 2 | 1 | Ampicillin-free plates | Focal | Strep_D2C1 | Illumina | ERS4256223 | No |
| +Com -Amp | 2 | 2 | Ampicillin-free plates | Focal | Strep_D2C2 | Illumina | ERS4256224 | No |
| +Com -Amp | 2 | 3 | Ampicillin-free plates | Focal | Strep_D2C3 | Illumina | ERS4256225 | No |
| +Com +Amp | 2 | 1 | Ampicillin-free plates | Focal | Strep_D2CA1 | Illumina | ERS4256226 | No |
| +Com +Amp | 2 | 2 | Ampicillin-free plates | Focal | Strep_D2CA2 | Illumina | ERS4256227 | No |

| | | | | | | | | |
|---|---|---|---|---|---|---|---|---|
| +Com +Amp | 2 | 3 | Ampicillin-free plates | Focal | Strep_D2CA3 | Illumina | ERS4256228 | No |
| -Com -Amp | 3 | 1 | Ampicillin-free plates | Focal | Strep_D3S1 | Illumina | ERS4256238 | No |
| -Com -Amp | 3 | 2 | Ampicillin-free plates | Focal | Strep_D3S2 | Illumina | ERS4256239 | No |
| -Com -Amp | 3 | 3 | Ampicillin-free plates | Focal | Strep_D3S3 | Illumina | ERS4256240 | No |
| -Com +Amp | 3 | 1 | Ampicillin-free plates | Focal | Strep_D3SA1 | Illumina | ERS4256241 | No |
| -Com +Amp | 3 | 2 | Ampicillin-free plates | Focal | Strep_D3SA2 | Illumina | ERS4256242 | No |
| -Com +Amp | 3 | 3 | Ampicillin-free plates | Focal | Strep_D3SA3 | Illumina | ERS4256243 | No |
| +Com -Amp | 3 | 1 | Ampicillin-free plates | Focal | Strep_D3C1 | Illumina | ERS4256235 | No |
| +Com -Amp | 3 | 2 | Ampicillin-free plates | Focal | Strep_D3C2 | Illumina | ERS4256236 | No |
| +Com -Amp | 3 | 3 | Ampicillin-free plates | Focal | Strep_D3C3 | Illumina | ERS4256237 | No |
| Basal +Amp | none | 1 | Ampicillin plates | Focal | Amp_BA1 | Illumina | ERS4256244 | Yes |
| Basal +Amp | none | 2 | Ampicillin plates | Focal | Amp_BA2 | Illumina | ERS4256245 | Yes |
| Basal +Amp | none | 3 | Ampicillin plates | Focal | Amp_BA3 | Illumina | ERS4256246 | Yes |
| +Com +Amp | 1 | 1 | Ampicillin plates | Focal | Amp_D1SA1 | Illumina | ERS4256247 | Yes |
| +Com +Amp | 1 | 2 | Ampicillin plates | Focal | Amp_D1SA2 | Illumina | ERS4256248 | Yes |
| +Com +Amp | 1 | 3 | Ampicillin plates | Focal | Amp_D1SA3 | Illumina | ERS4256249 | Yes |
| +Com +Amp | 3 | 1 | Ampicillin plates | Focal | Amp_D3SA1 | Illumina | ERS4256250 | Yes |
| +Com +Amp | 3 | 2 | Ampicillin plates | Focal | Amp_D3SA2 | Illumina | ERS4256251 | Yes |
| +Com -Amp | 1 | 1 | Ampicillin plates | Resident | Resident Ecoli D1C1 | Illumina | ERS4256252 | Yes |
| +Com -Amp | 1 | 2 | Ampicillin plates | Resident | Resident Ecoli D1C2 | Illumina | ERS4256253 | Yes |
| +Com -Amp | 1 | 3 | Ampicillin plates | Resident | Resident Ecoli D1C3 | Illumina | ERS4256254 | Yes |
| +Com +Amp | 1 | 1 | Ampicillin plates | Resident | Resident Ecoli D1CA1 | Illumina | ERS4256255 | Yes |
| +Com +Amp | 1 | 2 | Ampicillin plates | Resident | Resident Ecoli D1CA2 | Illumina | ERS4256256 | Yes |
| +Com +Amp | 1 | 3 | Ampicillin plates | Resident | Resident Ecoli D1CA3 | Illumina | ERS4256257 | Yes |
| +Com -Amp | 3 | 1 | Ampicillin plates | Resident | Resident Ecoli D3C1 | Illumina | ERS4256258 | Yes |
| +Com -Amp | 3 | 2 | Ampicillin plates | Resident | Resident Ecoli D3C2 | Illumina | ERS4256259 | Yes |
| +Com -Amp | 3 | 3 | Ampicillin plates | Resident | Resident Ecoli D3C3 | Illumina | ERS4256260 | Yes |
| +Com +Amp | 3 | 1 | Ampicillin plates | Resident | Resident Ecoli D3CA1 | Illumina | ERS4256261 | Yes |
| +Com +Amp | 3 | 2 | Ampicillin plates | Resident | Resident Ecoli D3CA2 | Illumina | ERS4256262 | Yes |
| +Com +Amp | 3 | 3 | Ampicillin plates | Resident | Resident Ecoli D3CA3 | Illumina | ERS4256263 | Yes |

| | | | | | | | | |
|---|---|---|---|---|---|---|---|---|
| +Com +Amp | 1 | 1 | Ampicillin plates | Resident | Resident Ecoli D1CA1 | MinION | ERS4256375 | Yes |
| +Com +Amp | 3 | 1 | Ampicillin plates | Resident | Resident Ecoli D3CA1 | MinION | ERS4256376 | Yes |
